# Structural Basis of Polypurine Track Strand Displacement by HIV-1 Reverse Transcriptase

**DOI:** 10.64898/2026.04.07.717013

**Authors:** Xin Wen, Rachel Lee, Sri Dhanya Muppalla, William M. McFadden, Karen A. Kirby, Robert A. Dick, Stefan G. Sarafianos

**Author notes:** Corresponding Author: Stefan G. Sarafianos.

## Abstract

To complete reverse transcription, HIV-1 reverse transcriptase (RT) must displace the RNase H-resistant polypurine tract (PPT) primers. This enables synthesis of the long terminal repeats and formation of the central cDNA flap. However, the molecular mechanism of this PPT strand displacement (SD) has remained unknown, and no structural data exist on how a retroviral polymerase execute these reactions. We report the first cryo-EM structures of HIV-1 RT bound to nucleic acid substrates containing either a PPT_RNA_ or PPT_DNA_ displacement strand, with incoming dATP positioned at the polymerase catalytic site. These structures reveal key features of the PPT displacement mechanism by RT. Specifically, we observed a binding mode where the template nucleotide (T_1_) base-paired to the first displacement nucleotide (D_1_) undergoes a 90° rotation relative to the preceding template base (T_0_). This sharp template flip positions D_1_ ∼30 Å away from the primer’s 3’-end and is coordinated by RT_p66_ residues at the SD interface: F61 and R78 contact T_1_/T_0_ to drive template translocation, while W24 engages both T_1_ and D_1_ to stabilize the displacement strand. Biochemical and virological mutagenesis experiments confirm that interactions with F61 and R78 are essential for both canonical cDNA polymerization and SD, whereas the W24-nucleotide interactions are required exclusively for SD but are dispensable for standard cDNA synthesis. These results contribute to the structural and functional understanding of PPT strand displacement by HIV-1 RT and reveal a distinct mechanistic vulnerability for the design of next-generation antiretrovirals.

## Introduction

Human Immunodeficiency Virus type 1 (HIV-1) reverse transcriptase (RT) is a multifunctional enzyme essential for converting the single-stranded viral RNA genome into double-stranded DNA (dsDNA) that is irreversibly integrated into the host genome^1–4^. HIV reverse transcription proceeds through a series of well-organized steps that requires precise coordination of the DNA polymerase and ribonuclease H (RNase H) activities of RT outlined in Figure 1^1,2^ (Fig. 1a-h). The process is initiated inside the viral capsid (CA) core after entry into the cytoplasm^1^. The host tRNA_Lys3_, packaged inside the CA core during viral assembly, binds to the primer binding site (PBS) on the viral RNA (Fig. 1a); RT then synthesizes the minus-strand strong-stop [(−)ssSTOP] cDNA at the 5’-end of the genome^5^ (Fig. 1b). Concurrently, the RNase H subdomain degrades the RNA template within the RNA/DNA hybrid, thus facilitating the first strand transfer of the (−)ssSTOP cDNA to the 3’ repeat (R) region of the viral genome, generating the first-strand transfer (FST) intermediate^6,7^ (Fig. 1c). RT continues (−)cDNA synthesis, generating the first U3-R-U5 sequence known as the long terminal repeat (LTR; Fig. 1d), while RNase H degrades most of the viral RNA template that generates the full-length minus-strand (FLM) intermediate^6,7^ (Fig. 1e).

**Figure 1:**
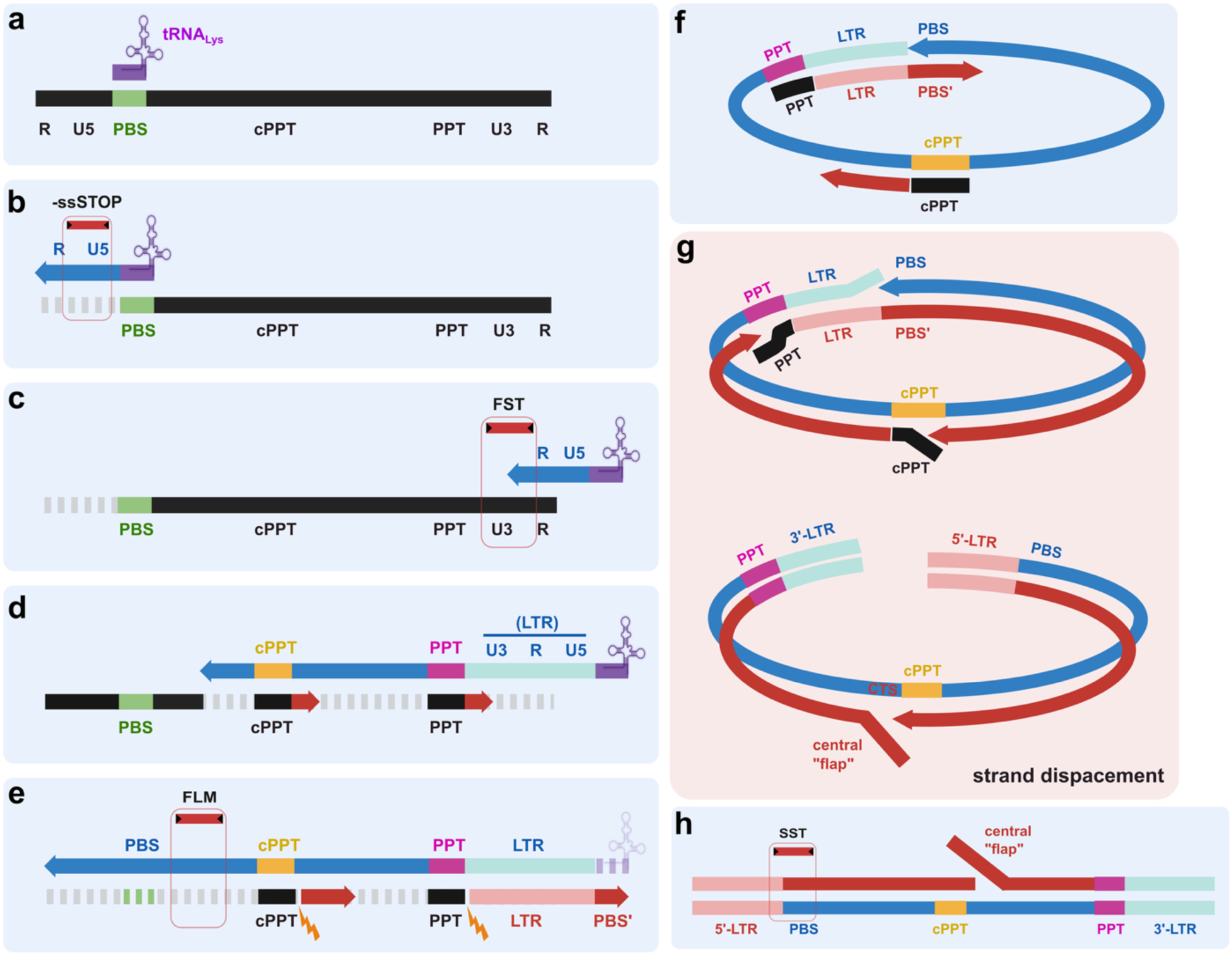
HIV-1 reverse transcription. **a** Host tRNA_Lys3_(purple) binds to the primer binding site (PBS, green) of the viral RNA genome to serve as a primer. **b** Synthesis of (−)cDNA initiates toward the 5’-end of the RNA genome, generating the -ssSTOP product. RNase H degrades the RNA template within the resulting RNA/DNA hybrid. **c** The first “jump” is facilitated by homologous annealing of the R regions, yielding the first strand transfer (FST) product upon further cDNA synthesis. **d** As (−)cDNA synthesis continues, it forms the U3-R-U5 sequence, known as the long terminal repeat (LTR). Meanwhile, RNase H degrades the RNA genome but skips the 3’-polypurine tract (PPT, magenta) and central PPT (cPPT, yellow), which serve as primers for (+)cDNA synthesis. **e** Elongation of the (−)cDNA strand continues, producing the full-length minus-strand (FLM) product. Following (+)cDNA synthesis initiation, RNase H cleaves the borders of the PPTs, detaching them from the newly synthesized (+)cDNA. The anti-PBS (PBS’) is then synthesized on the (+)cDNA, followed by tRNA_Lys3_ removal by RNase H. **f** The intramolecular second strand transfer (second “jump”) is driven by homologous annealing of the PBS regions, forming a circular cDNA intermediate. **g** Strand displacement occurs at the 3’-ends of the U5-R-U3 region, the 3’-PPT, and the cPPT. This resolves the circular intermediate and completes the copying of the two LTRs. Additionally, a central “flap” is generated at the central termination sequence (CTS). **h** The resulting full-length, linearized proviral dsDNA is ready for integration into the host genome. Black: viral RNA genome; Blue: (−)cDNA; Red: (+)cDNA. The 5’-LTR and 3’-LTR are indicated by pink and light blue lines, respectively. Strand displacement steps are grouped within the large pink box. Red boxes denote early (-ssSTOP and FST), intermediate (FLM), and late (SST) reverse transcription products. The red lines with triangles indicate qPCR primer targeting sites for detecting these corresponding products. Schematics created in BioRender. Wen, X. (2026) https://BioRender.com/vfsz9ra.

RT RNase H cleavage of the HIV-1 genome is not uniform throughout the nucleic acid strand^8,9^. Specific purine-rich sequences known as polypurine tracts (PPTs) are relatively resistant to RNase H cleavage and remain annealed to the nascent (−)cDNA^8–10^ (Fig. 1d). The uncleaved 3’-PPT and central PPT (cPPT) that are identical sequences serve as primers for (+)cDNA synthesis^11–16^ (Fig. 1d). Although it has been suggested that certain PPT-like RNA fragments could also remain bound to the viral cDNA and serve as primers, these events are likely less dominant in HIV-1 reverse transcription compared to PPT-primed synthesis^6,17,18^. Thus, the predominant primers are the 3’-PPT, which initiates synthesis of the upstream (+)cDNA to the LTR region, and the cPPT, which initiates synthesis of the middle segment^11,14,15,19^ (Fig. 1d). Following (+)cDNA initiation, RNase H cleaves the RNA/DNA junctions of the PPT primers^14,20,21^, potentially leaving the PPTs bound to the (−)cDNA as (+)cDNA synthesis continues along the strand (Fig. 1e). RNase H also removes the tRNA primer leaving behind a single rA nucleotide at the 5’ end of the (−)cDNA^14^ (Fig. 1e). This removal enables the intramolecular second strand transfer where the (−)cDNA PBS region anneals to the (+)cDNA PBS’, forming a circular cDNA intermediate^1,6,7^ (Fig. 1f).

To resolve this circular intermediate and generate a linear, integration-competent dsDNA with an LTR at each end, RT must facilitate proper strand displacement (SD) ^22–27^ (Fig. 1g). It has been proposed that (−)cDNA synthesis is complete after displacement of the U5, R, U3 sequences at the 5’-end, thus the circular intermediate is opened and the 5’-LTR is copied^22,23,28^ (Fig. 1g). Successful removal of the PPTs is also essential for viral replication^29^; therefore, it is likely that RT also displaces the 3’-PPT_RNA_ primer that was annealed to the (−)cDNA and finishes copying the 3’-LTR (Fig. 1g). Additionally, the 3’-PPT-primed (+)cDNA synthesis is eventually elongated to the residual cPPT_RNA_ primer (Fig. 1g). At this junction, RT displaces the cPPT and continues SD synthesis ∼100 nucleotides (nt) into the (+)cDNA until it reaches the central termination sequence (CTS), generating a central DNA “flap” essential for nuclear import and viral replication in a cell type-dependent manner ^26–33^ (Fig. 1g, bottom). The linearized dsDNA is ready for host-genome integration (Fig. 1h).

While HIV-1 RT possesses the intrinsic ability to displace extensive downstream nucleic acids during cDNA synthesis^22–25^, the displacement of the PPT_RNA_ involves substrates that are unique in their structural characteristics^10,34–36^. Unlike a standard sequence, PPTs comprise consecutive purine ribonucleosides and form RNA/DNA hybrids with the DNA template^10,19–21^. Structural and biophysical studies have demonstrated that polypurine hybrids are generally more rigid and can adopt a conformation distinct from standard B-form DNA^10,35–38^. Crystal structures of the HIV-1 RT bound to a PPT RNA/DNA indicate that these sequences can undergo a structural “slippage”^10^, which has been observed to result in out-of-register base pairing and mismatched nucleotides that can alter the physical properties of the nucleic acid hybrid^10,16,38^. Successfully displacing these PPT_RNA_ primers is critical for viral genomic integrity and host-genome integration^30,39^. At the 3’-end, failure to fully displace the PPT prevents the formation of complete LTRs, resulting in aberrant “1-LTR circles“^4,40^. These circles are incapable of integration, though recent studies indicate they are capable of sustaining some level of transcription^40–42^. At the center of the genome, cPPT-initiated synthesis reduces the temporal exposure of single-stranded (−)cDNA, minimizing the window for lethal editing by host restriction factors such as APOBEC3G/F^43,44^. Biochemical and mutational reports have implicated RT residues that affect SD^45–47^. Despite the numerous reports on these events, the molecular mechanism by which RT performs PPT-based SD remains poorly understood.

The structure of HIV-1 RT has been extensively characterized by X-ray crystallography and cryo-electron microscopy (cryo-EM) across various functional states^48–50^. High-resolution structures of elongation complexes with DNA/RNA or DNA/DNA have revealed the pre- and post-translocation states of RT^51,52^. Furthermore, ternary complexes capturing incoming dNTPs have elucidated the catalytic cycle of nucleotide addition^52–55^. Recent cryo-EM structures have captured HIV-1 reverse transcription initiation complexes and dynamic dATP incorporation intermediates, providing snapshots of the enzyme’s conformational flexibility^56,57^. In the broader field of DNA polymerases, recent structures focused on DNA strand displacement have been reported^58,59^. For the yeast mitochondrial DNA polymerase Mip1, cryo-EM structures revealed a “wedge helix” and a “catcher loop” that coordinate to actively separate the downstream DNA/DNA duplex^58^. Additionally, a recently reported cryo-EM structure of human Pol γ was proposed to have a passive mechanism where the enzyme does not actively displace the strands, but rather it captured four different conformational states of the spontaneously unwinding downstream DNA duplex^59^. To date, there is no structural information describing PPT-containing strand-displacement complexes, including PPT_RNA_/DNA or PPT_DNA_/DNA. Moreover, there is no structural information on the molecular determinants of how retroviral polymerases engage in specific strand displacement reaction, a process essential for the completion of genome replication.

To bridge this gap and better understand the molecular basis of PPT strand displacement by HIV-1 RT, we determined structures in complex with displacement PPT-based nucleic acid substrates and an incoming dATP using single-particle cryo-EM. We report structures with either a PPT_DNA_ displacement strand or the physiological PPT_RNA_ displacement strand, allowing us to examine potential mechanistic distinctions between DNA and RNA strand displacement by RT. We also report the role of distinct helical properties between the DNA/DNA duplex and the RNA/DNA hybrid, resulting in differential substrate interactions with residue T290 in the p66 thumb subdomain. Our structures revealed p66 residues W24, F61, and R78 interacting with the nucleic acid substrate at the SD site, likely facilitating the displacement duplex separation. Biochemical and virological assays confirmed that W24 critically and specifically affects the SD activity of RT without significantly impacting normal polymerization, whereas F61 and R78 play important roles on standard DNA synthesis, thereby also impacting strand displacement.

## Results

### Cryo-EM structure of HIV-1 RT in complex with PPT_DNA_ displacement substrate and incoming dATP (RT/T-P-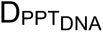/dATP)

We first determined the cryo-EM structure of the RT/T-P-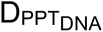/dATP ternary complex, where the downstream displacement starts with a PPT_DNA_ (Fig. 2, Supplementary Figs. 1-2). In this 2.6 Å resolution structure, the primer strand is covalently crosslinked to the RT_p66_ thumb subdomain, and the PPT_DNA_ displacement strand is derived from the HIV-1 CTS containing the cPPT (Fig. 2a). The resulting cryo-EM map reveals a previously unobserved density extending upward between the p66 fingers and thumb subdomains, which we interpret as the PPT_DNA_ displacement duplex (Fig. 2b-c). The local resolution near the enzyme active site, the template-primer (T-P), and the first base-pairs of the displacement duplex is ≤3 Å (Supplementary Fig. 2a-b). Resolution of the displacement duplex decreases towards the distal end due to flexibility (from ∼3 to ∼6 Å; Supplementary Fig. 2a-b). Consequently, we confidently modeled the RT/T-P and incoming dATP but limited the initial model of the displacement duplex to the first 10 base-pair (bp) (Fig. 2c). To improve the local density quality of the displacement duplex, we performed 3D auto-refinement using an extended mask covering the downstream region, yielding a 2.7 Å map (Supplementary Fig. 1, Supplementary Fig. 2a-c). Using a Gaussian lowpass-filtered map and strong secondary-structure constraints during model refinement allowed us to extend the PPT_DNA_ model to 15 bp, covering the full cPPT sequence. (Supplementary Fig. 2c).

**Figure 2:**
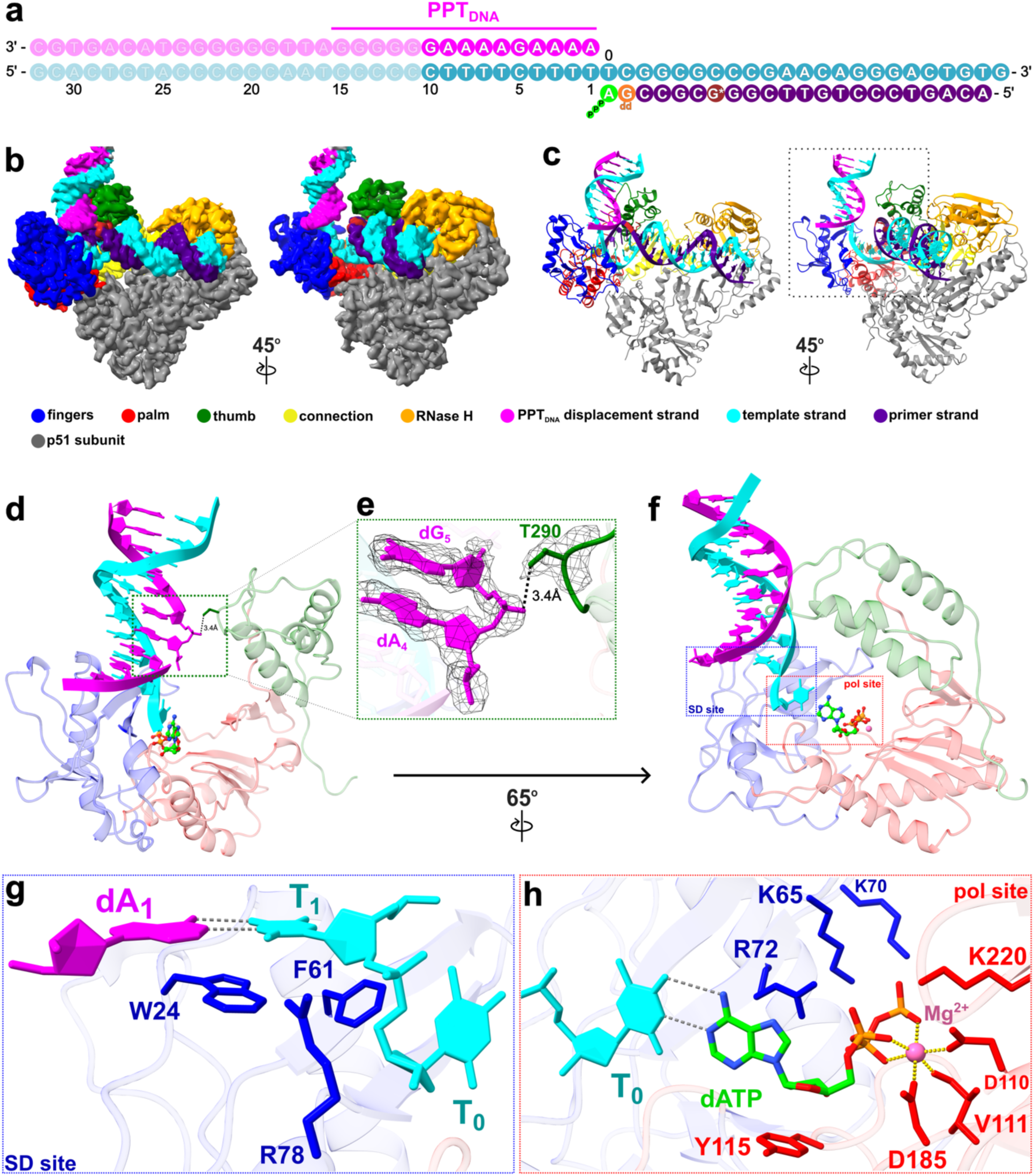
Cryo-EM structure of HIV-1 RT in complex with PPT_DNA_ displacement substrate and incoming dATP (RT/T-P_ddG_-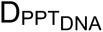/dATP). **a** Schematic of the nucleic acid substrate with the primer strand in purple, the template strand in cyan, and the displacement strand in magenta (PPT_DNA_ residues are indicated by a bar). Modifications on primer strand include a thioalkyl-tethered guanosine (G*, maroon) and a dideoxy guanosine (ddG, orange). The incoming deoxyadenosine triphosphate (dATP) is in green. Faded regions of the nucleic acid indicate segments not modeled in the structure. **b** Cryo-EM density map and **c** ribbon model are shown in front (left) and 45° rotated side views (right). The map is contoured at 6.7σ. RT_p66_ is colored by subdomain: fingers (blue), palm (red), thumb (green), connection (yellow), and RNase H (orange). RT_p51_ is shown in gray. **d** Close-up view of the polymerase domain (boxed region in **c**), showing the displacement duplex and incoming dATP (lime green, atom-colored). **e** Detailed view of the interaction between T290 and the displacement strand (boxed region in **d**). The cryo-EM density (mesh) is from the postprocess map contoured at 5.8σ. The distance between the T290 Cɣ2 atom and the dG5 phosphate oxygen is indicated. **f** Structural overview rotated 65° relative to **d**, highlighting the relative position of the strand displacement (SD) site (blue box) and the polymerase (pol) active site (red box). **g** Detailed view of the SD site (**blue box in f**) showing the primary interacting residues W24, F61, and R78 in vicinity of the first displacement base pair, dA_1_-T_1_. **h** Detailed view of the pol site (**red box in f**) showing key residues involved in dATP binding. The Mg^2+^ ion (pink sphere) is coordinated (yellow dash lines) by six surrounding oxygen atoms. Dashed gray lines indicate hydrogen bonds.

The unexpected positioning of the PPT_DNA_ duplex between the p66 fingers and thumb revealed novel interactions at the polymerase subdomain (Fig. 2d). Residue T290 of the p66 thumb lies in proximity to the displacement strand and appears to form a weak interaction with the phosphate backbone between the 4th and 5th nucleotides (dA_4_-dG_5_), potentially stabilizing the upward twist of the displacement duplex (Fig. 2e). Rotating the structure view by 90° highlights the SD site and the polymerase active site simultaneously (Fig. 2f). As expected, the incoming dATP pairs with the last template nucleotide before the displacement region (T_0_). Surprisingly, the subsequent template nucleotide (T_1_), which pairs with the first nucleotide of the DNA displacement strand (dA_1_), is rotated by ∼90° relative to T_0_. As a result, the displacement strand assumes an unusual conformation, with the 5’-end of the PPT_DNA_ located ∼33 Å away from the 3’-OH of the primer terminus at the polymerase active site (Supplementary Fig. 2d).

This unusual conformation appears to be stabilized by interactions of the template-displacement (T-D) duplex with aromatic residues W24 and F61 at the SD site (Fig. 2g). While F61 interacts with T_1_ and T_0_, W24 stacks with both T_1_ and dA_1_ (Fig. 2g; Supplementary Fig. 2e-h). Moreover, R78 interacts with the phosphate backbone between T_0_ and T_1_ (Fig. 2g; Supplementary Fig. 2e-h), potentially steering template translocation to facilitate strand displacement. Finally, at the polymerase active site, the side chains of residues K65, K70, R72, D110, V111, Y115, D185, and K220 coordinate the incoming dATP with the Mg^2+^ ions as described in previous structures^6,48,49,51,57^ (Fig. 2h; Supplementary Fig. 2g).

### Structural comparison of catalytic displacement complexes of RT with PPT_DNA_ and PPT_RNA_: RT/T-P-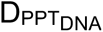/dATP and RT/T-P-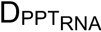/dATP

To examine if these SD features persist with the native cPPT_RNA_ substrate, we determined the cryo-EM ternary structure of RT/T-P-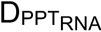/dATP at 3 Å resolution (Supplementary Fig. 3). The experimental workflow followed that of the PPT_DNA_ complex but featured a displacement strand starting with the PPT_RNA_ (Fig. 3a, Supplementary Fig. 4a). As observed previously, local resolution is highest over the RT and T-P, while the displacement region varies from ∼3 - 8 Å (Supplementary Fig. 4a-b). We confidently modeled the first 10 bp of the RNA-displacement/DNA-template duplex (Fig. 3b). Using the same approach described previously, we extended the model to 17 bp, capturing two DNA base pairs beyond the PPT_RNA_ (Supplementary Fig. 4a-c).

**Figure 3:**
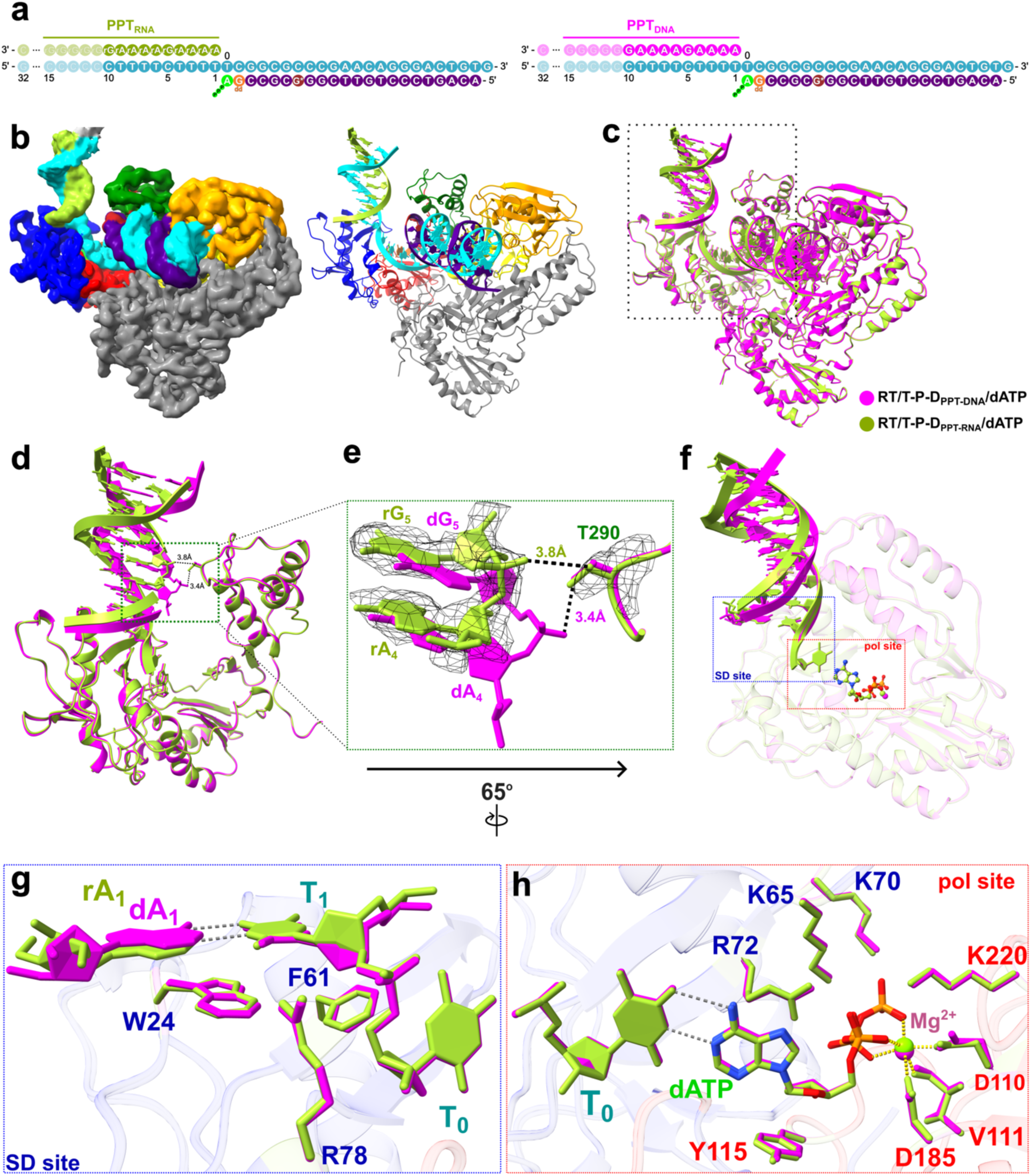
Structural comparison of RT-PPT_DNA_ and RT-PPT_RNA_ displacement complexes: RT/T-P-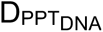/dATP and RT/T-P-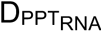/dATP. **a** Schematic of the nucleic acid substrates containing either a PPT_RNA_ (left) or PPT_DNA_ (right) displacement strand. The primer strand (purple) contains modifications (ddG, G*) colored as in Fig. 2. The template strand is cyan, and the incoming dATP is green. The displacement strand is shown in magenta (PPT_DNA_) or yellow-green (PPT_RNA_). Faded regions of the nucleic acids indicate segments not modeled. **b** Cryo-EM density map (left) and ribbon model (right) of the RT-PPT_RNA_ displacement complex. Density map is contoured at 7.4σ. RT_p66_ and RT_p51_ subdomains are colored as in Fig. 2. **c** Superposition of the RT-PPT_DNA_ (magenta) and RT-PPT_RNA_ (yellow-green) complexes. The structures align closely at the polymerase active site and primer-template region (RMSD 0.74 Å) but diverge significantly at the downstream displacement duplex (RMSD 4.99 Å). **d** Close-up view of the polymerase domain (boxed region in **c**). **e** Detailed view of the T290 interaction (boxed region in **d**). The PPT_RNA_ complex density (mesh) is from the postprocess map contoured at 5.0σ. Dashed lines indicate the distance between the T290 side chain and the displacement strand phosphate for DNA (3.4 Å) versus RNA (3.8 Å). **f** Structural overview rotated 65° relative to **d**, highlighting the distinct locations of the strand displacement (SD) site (blue box) and polymerase (pol) active site (red box). **g** Detailed view of the SD site (**blue box in f**). Positions of key interacting residues (W24, F61, R78) and nucleic acids show high structural similarity. **h** Detailed view of the pol active site (**red box in f**). Labeled residues, incoming dATP, and Mg^2+^ align closely.

Overall, the PPT_DNA_ and PPT_RNA_ complexes show high structural similarity, with near-perfect overlap in the enzyme and T-P regions (Fig. 3c, 3g-h; Supplementary Fig. 4d-g). The key differences are found at the displacement duplex (Fig. 3d-f), consistent with the distinct helical geometry of RNA-DNA and DNA-DNA duplexes. The distinct helical geometry appears to reposition the RNA-DNA displacement duplex, affecting the distance between the PPT_RNA_ and RT-T290 (Fig. 3e).

As the displacement duplex is highly flexible in both complexes, the apparent T290 contact in the consensus maps likely reflects an average over multiple conformations rather than a fixed interaction. To quantify this flexibility, we classified the final particle stack from each dataset into five 3D classes to capture distinct conformations of the downstream displacement duplex (Supplementary Fig. 5a-h). In both datasets, the five classes superimpose closely at the enzyme region but vary in the displacement duplexes (Supplementary Fig. 5b-d, f-h). Focusing first on the PPT_DNA_/p66-thumb interface, our analysis revealed that ∼59% of particles show a weak interaction at 3.4-3.5 Å between T290 and the PPT_DNA_ displacement strand, while the remaining ∼41% of the particles show a loss of interaction at 3.6-3.8 Å distances (Supplementary Fig. 5d). The PPT_RNA_ dataset demonstrates a similar variability in the movement of the displacement duplex (Supplementary Fig. 5e-h). Here, ∼44% of particles retain the weak interaction (3.3-3.4 Å distance), whereas the remaining ∼56% show a larger separation of 3.8 Å (Supplementary Fig. 5h). Consistent with the 3D classification analyses, 3D Variability Analysis (3DVA) of both datasets displays the PPT_DNA_ and PPT_RNA_ duplexes swaying side-to-side, contacting the p66 thumb only in a subset of conformations (Supplementary Videos 1-2). These structural analyses reveal that the PPT-T290 interaction is modest and a dynamic feature common to both PPT_DNA_ and PPT_RNA_ substrates.

Despite the structural differences of the displacement duplexes, the two complexes superimpose closely at the SD and polymerase active sites (Fig. 3f, Supplementary Fig. 4e-g). The first displacement-template base pairs, rA_1_-T_1_ and dA_1_-T_1_, align well, and the contacts mediated by W24, F61, and R78 are similar in the PPT_RNA_ and PPT_DNA_ complexes (Fig. 3g). The polymerase site geometry and the coordination of the incoming dATP remain highly similar in both complexes (Fig. 3h). Collectively, these observations suggest that W24, F61, R78, and T290 may play important roles in HIV-1 RT strand displacement.

### Biochemical validation of structural interactions at the strand displacement site

To validate the functional impact of W24, F61, R78, and T290 on RT strand displacement, we generated a targeted panel of 13 RT mutants. This panel includes “loss-of-function” mutants (W24A, F61A, R78A, T290A/D) and “conservative” mutants (W24F, F61L, R78K, T290S). Additionally, we constructed double mutants targeting the W24-F61 pair, as these residues are positioned directly beneath the first T-D base pair.

#### Primer extension on a T-P-only substrate

We first assessed the intrinsic polymerization activity of these mutants using a standard T-P substrate that does not involve strand displacement^60^ (Fig. 4a, Supplementary Fig. 6a). As expected, RT-WT extends primers to full length in both 15-and 60-min reactions (Fig. 4b, Supplementary Fig. 6b-d). W24A and W24F exhibit polymerization activity comparable to WT at both time points (Supplementary Fig. 6b). While F61A shows a modest reduction in primer extension at 15 min (Fig. 4b), it reaches full-length extension by 60 min, and the conservative F61L mutant displays near-WT activity throughout (Supplementary Fig. 6b). Among the double mutants, W24A/F61A exhibits a strong defect in polymerization^47^, with consistently reduced full-length product, whereas the conservative double mutants retain WT-level activity (Fig. 4b, Supplementary Fig. 6c). Finally, all R78 and T290 mutants retain WT-level polymerization activity (Fig. 4b, Supplementary Fig. 6d).

**Figure 4:**
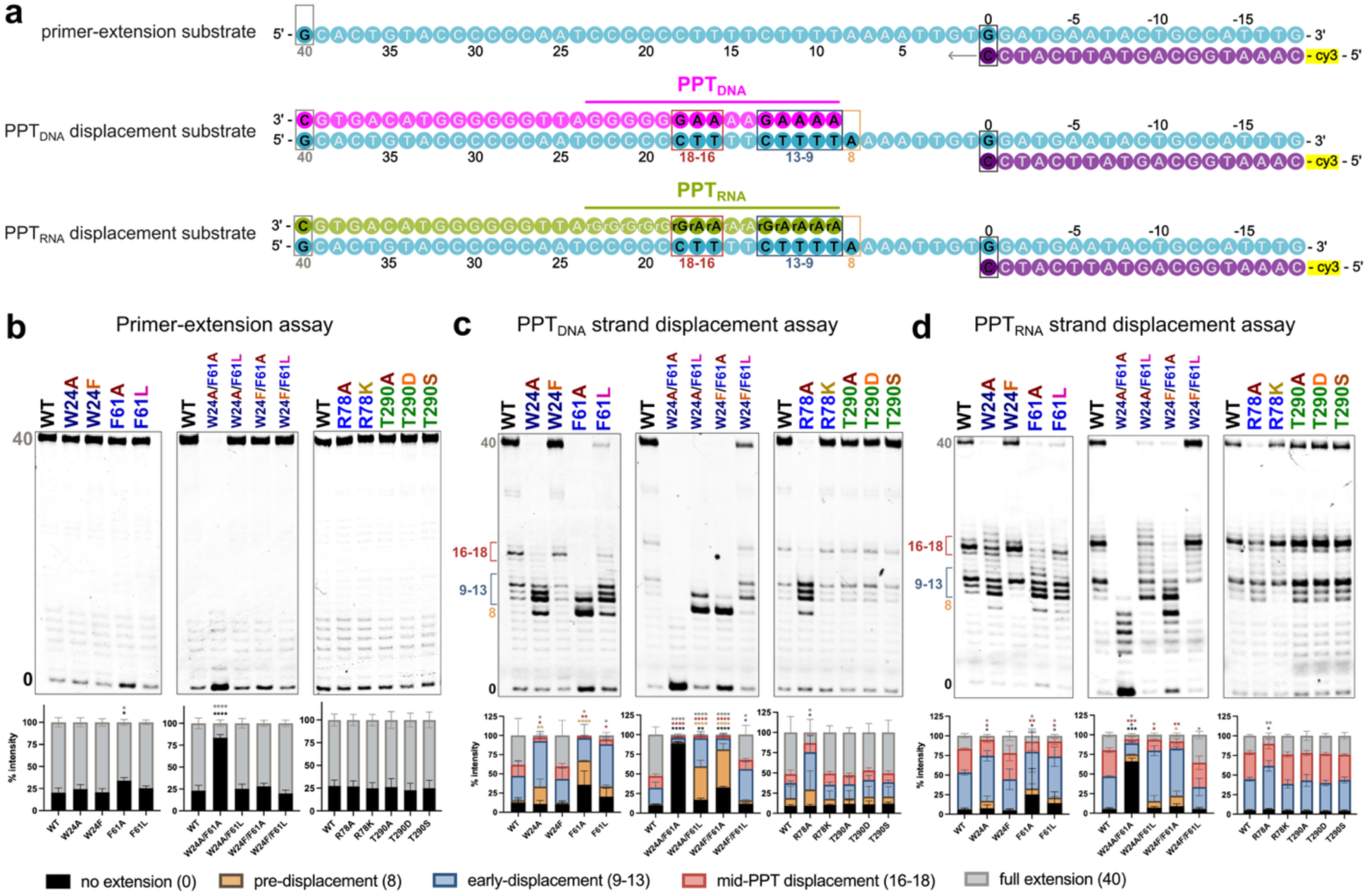
Impact of HIV RT strand-displacement (SD) site mutations on the primer-extension and SD activities. **a** Schematics of the nucleic acid substrates used in the biochemistry assays: the extension-only template-primer substrate (top), the extension and PPT_DNA_ displacement substrate (middle), and the extension and PPT_RNA_ displacement substrate (bottom). The primer strand (purple) is labeled with a Cy3 fluorophore at the 5’-end. The template strand is cyan, the PPT_DNA_ displacement strand is magenta, and the PPT-RNA displacement strand is yellow-green. Boxed, darker-colored nucleotides represent strong primer-extension pause sites observed in the assays. **b-d** Representative PAGE gels and corresponding quantitative bar graphs for primer-extension assays (**b**) PPT_DNA_ strand displacement assay (**c**), and PPT_RNA_ strand displacement assay (**d**) with RT-WT and the indicated mutants. Gel bands represent the size of the Cy3-labeled primer after 15-minute reactions at a 2:1 RT:substrate ratio (**b, c**) or 30-minute reactions at a 4:1 RT:substrate ratio (**d**). Key pausing-positions of interest are defined and color-coded as follows: position **0**, un-extended primer (black); position **8**, primer extension stopped at the pre-displacement site (orange); positions **9-13**, primer stopped at the early-displacement site (blue); positions **16-18**, primer stopped at the mid-PPT displacement site (red); and position **40**, fully extended primer (gray). This color coding is consistently applied to the stacked bar graphs below the gels, which represent the quantification of these bands from three independent experiments. Additional time points and triplicate gels are shown in **Supplementary figures 6-8**. Statistical significance was determined using a one-way ANOVA with Dunnett’s multiple comparisons test, comparing each mutant to the WT control on the corresponding gel. *P < 0.05; **P < 0.01; ***P < 0.001; ****P < 0.0001; bars without asterisks are not significant (P > 0.05).

#### Strand displacement on T-P-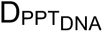

To evaluate the impact of these mutants on the SD activity of RT, we performed the same assay with nucleic acid substrate containing a downstream PPT_DNA_ displacement strand (Fig. 4a, Supplementary Fig. 7a). RT-WT efficiently extends >50% of primers to the end of the template within 15 min and achieves near-complete primer extension by 60 min (Fig. 4c, Supplementary Fig. 7b-d). We observe some pausing at the PPT region at early time points, which resolves over time, indicating efficient strand displacement by RT-WT (Fig. 4c, Supplementary Fig. 7b-d). W24A extends primers normally up to the SD site but is strongly impaired in strand displacement activity (Fig. 4c); at 15 min, primers stall at pre- and early-displacement positions, and only ∼30% reach full length after 60 min (Supplementary Fig. 7b). This defect is rescued in W24F (Fig. 4c, Supplementary Fig. 7b), indicating that an aromatic side chain at position 24 is critical for strand displacement. While F61A reduces polymerization at 15 min (Fig. 4c), at later time points most primers initiate extensions but predominantly stall at pre- and early-displacement sites (Supplementary Fig. 7b). Despite the improvement in overall extension over time, only ∼20% of primers reach full length by 60 min (Supplementary Fig. 7b). F61L partially rescues this defect but does not fully restore WT-level activity, validating the importance of the aromatic character of F61 (Fig. 4c, Supplementary Fig. 7b).

Consistent with the cooperative structural positioning of residues 24 and 61, the W24A/F61A double mutant is severely defective in both polymerization and strand displacement, with most primers remaining un-extended even after 60 min (Fig. 4c, Supplementary Fig. 7c). The extended primers fail to even reach the displacement site (Fig. 4c). The single-rescue mutants, W24A/F61L and W24F/F61A, partially restore polymerization but cannot support strand displacement at 15 min (Fig. 4c). By 60 min, while most primers have initiated extension, they predominantly stall at pre- and early-displacement sites (Supplementary Fig. 7c). The double-rescue mutant, W24F/F61L, fully restores both activities to near-WT levels (Fig. 4c, Supplementary Fig. 7c), highlighting the importance of coordinated hydrophobic/aromatic interactions at the SD site. Furthermore, R78A shows reduced strand displacement with stalling at pre- and early-displacement sites, while R78K restores WT-level activity (Fig. 4c, Supplementary Fig. 7d). Lastly, although a weak transient contact between T290 and the PPT_DNA_ backbone is observed in the consensus EM map, mutations T290A, T290D, and T290S have no significant effect on polymerization nor strand displacement (Fig. 4c, Supplementary Fig. 7d). This is consistent with our structural analysis indicating that the T290 contact is present only in a subset of particle conformations (Supplementary Fig. 5).

#### Strand displacement on T-P-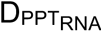

We also examined strand displacement using the physiological displacement substrate of HIV-1 RT, PPT ^22^ (Fig. 4a, Supplementary Fig. 8a). Overall, RT-WT displays reduced strand displacement efficiency on PPT_RNA_ compared to PPT_DNA_ (Fig. 4c-d, Supplementary Fig. 7-8). Even with increased enzyme concentration and extended reaction times, we observe substantial pausing at early- and mid-displacement sites, with only ∼30% of primers fully extended after 90 min (Supplementary Fig. 8b-d). Despite the reduced efficiency, the relative effects of the RT mutations are similar to those observed with the PPT_DNA_. W24A and F61A remain defective in strand displacement, with primers stalling at pre- and early-displacement sites, whereas W24F fully rescues activity to WT levels and F61L provides partial rescue (Fig. 4d, Supplementary Fig. 8b). W24A/F61A shows minimal primer extension into the displacement region at 30 min (Fig. 4d); by 90 min, approximately half of the primers reach and stall at the pre- and early-displacement sites (Supplementary Fig. 8c). Strand displacement is partially rescued by W24A/F61L and W24F/F61A, while W24F/F61L restores activity to near-WT levels (Fig. 4d, Supplementary Fig. 8c). R78A shows reduced PPT_RNA_ strand displacement that is rescued by R78K, while all T290 mutants behave comparably to RT-WT (Fig. 4d, Supplementary Fig. 8d). Together, these results confirm that W24, F61, and R78 play important roles in RT strand displacement across both PPT_DNA_ and PPT_RNA_ substrates, whereas T290 does not substantially contribute to this activity in these biochemical assays.

### Cellular and viral validation of RT strand displacement mutants

To validate the biological relevance of W24, F61, R78, and T290, we introduced these mutations into the pNL4-3ΔEnv construct and generated VSV-G-pseudotyped HIV-1 virions for single-cycle infection assays in TZM-GFP cells^61^ (Fig. 5a). All mutant viruses were normalized to p24 concentration prior to infection (Supplementary Fig. 9a). Consistent with our biochemical data, loss-of-function single mutants (W24A, F61A, R78A) and the double mutant (W24A/F61A) display a complete loss of infectivity (Fig. 5b). Similarly, single-rescue double mutants (W24A/F61L and W24F/F61A) exhibit no detectable infectivity (Fig. 5b), confirming that restoring only one residue of the hydrophobic pair at the SD site is insufficient for viral replication. The conservative mutants W24F and F61L fully restore infectivity to WT levels (Fig. 5b). The R78K and the double-rescue W24F/F61L mutants restore infection efficiency to ∼75% of WT (Fig. 5b). Interestingly, while T290 mutants generally maintain WT-level infectivity (Fig. 5b), the T290D mutant exhibits a slight but significant reduction compared to T290A (Supplementary Fig. 9b), suggesting a minor contribution to optimal viral replication.

**Figure 5:**
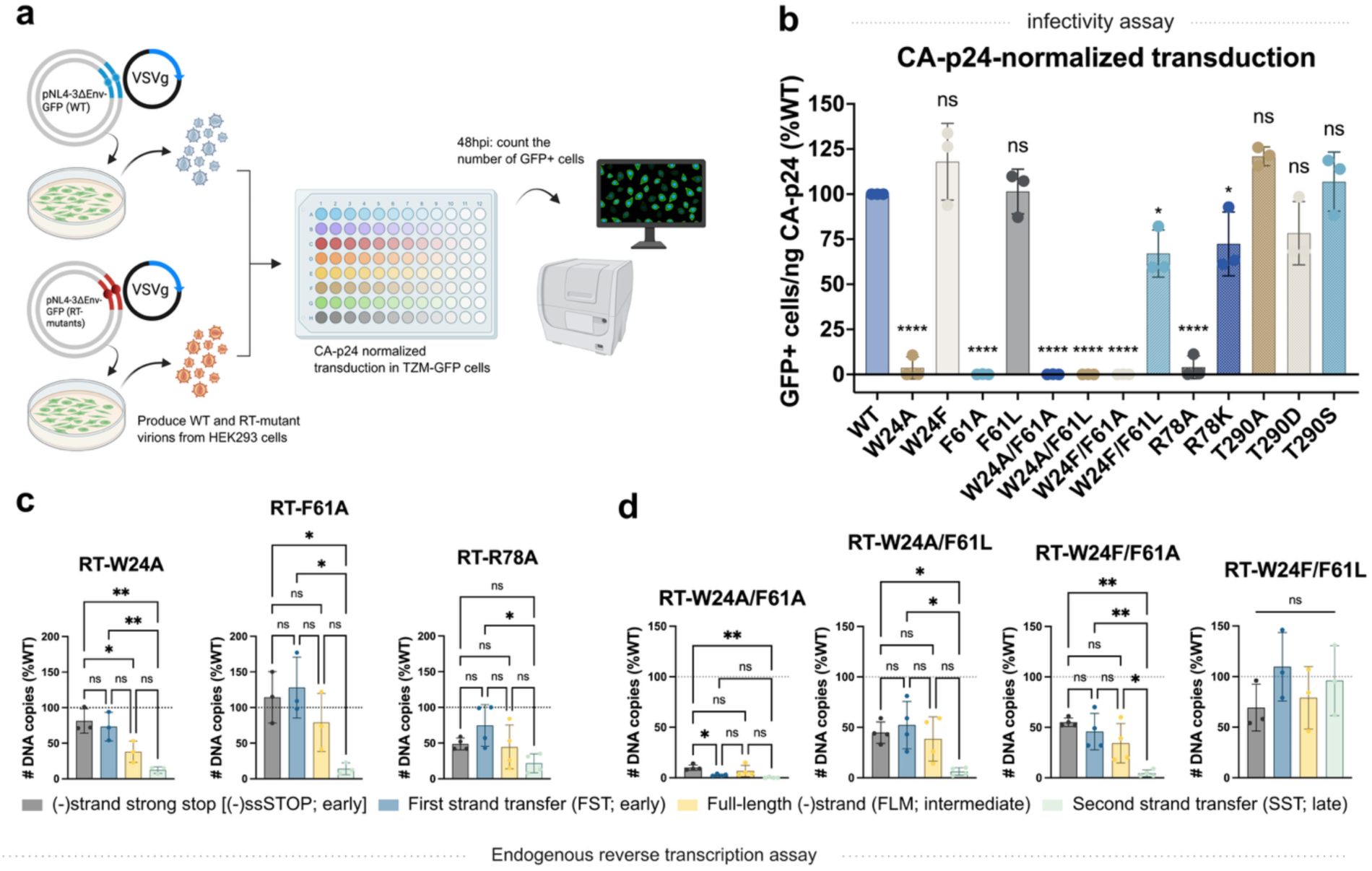
Virological impact of strand-displacement site mutations on HIV-1 infection and viral cDNA synthesis. **a** Schematic of the pseudotyped HIV-1 (WT and RT mutant) transduction assay. **b** CA-p24-normalized infectivity of pseudotyped HIV-1 virions (WT and RT mutants) in TZM-GFP cells. The number of GFP+ cells was measured at 48 hpi and normalized to the WT. Bar graphs show the mean of three independent experiments. Statistical significance was determined using a one-way ANOVA with Dunnett’s multiple comparisons test, comparing each mutant to the WT control. *P < 0.05; ****P < 0.0001; ns, not significant. **c-d** Endogenous reverse transcription (ERT) assays of single-mutant (**c**) and double-mutant (**d**) virions that exhibited a complete loss of infectivity in **b**. Bar graphs show the number of DNA copies for products at different stages of reverse transcription as indicated in Fig. 1 ((−)strand strong stop, first strand transfer, full-length (−)strand, and second strand transfer), normalized to the corresponding product level in WT virions. Bar graphs show the mean of three or four independent experiments. Statistical significance was determined using a one-way ANOVA with Tukey’s multiple comparisons test, comparing the relative levels of each RT product to one another. *P < 0.05; **P < 0.01; ns, not significant. Schematics created in BioRender. Wen, X. (2026) https://BioRender.com/ioa1i5f.

To better understand the impact of these mutants on specific steps of the reverse transcription process at the viral level, we employed an endogenous reverse transcription (ERT) assay^62,63^. This method allows quantification of reverse transcription intermediates synthesized inside native viral capsid cores but is isolated from the cellular environment. We monitored the synthesis of early- (-ssSTOP and FST), intermediate- (FLM), and late-stage (SST) products (Fig. 1). W24A and F61A synthesize early and intermediate products at levels comparable to WT but show a severe, defect in synthesizing late-stage product (Fig. 5c). As the synthesis of late-stage products directly involves the PPT strand displacement step (Fig. 1g-h), this specific reduction implicates a failure in displacement activity. This reduction is fully restored in the W24F and F61L viruses (Supplementary Fig. 9c).

R78A reduces the early- and intermediate-products by ∼50%, and the late-product to ∼30% of WT (Fig. 5c), indicating impairments in both general polymerization and strand displacement functions. R78K rescues all synthesis steps, though not fully (Supplementary Fig. 9c). Consistent with the biochemical data, W24A/F61A fails to generate even early reverse transcription products (Fig. 5d), confirming a total loss of polymerase function. While the single-rescue double mutants (W24A/F61L and W24F/F61A) partially restore the synthesis of early and intermediate products, they fail to synthesize detectable levels of late-stage products (Fig. 5d), highlighting that the W24-F61 pair is essential for strand displacement activity. The double-rescue mutant W24F/F61L fully restores synthesis of all viral cDNA intermediates (Fig. 5d). Finally, viruses carrying T290 mutations show no significant difference between reverse transcription stages within each virus (Supplementary Fig. 9d). However, overall product levels vary: T290A and T290D mutants display a slight but consistent reduction in all cDNA intermediates compared to WT, whereas T290S exhibits levels exceeding 100% of WT (Supplementary Fig. 9d). These results suggest that T290 may mildly impact the overall efficiency of reverse transcription.

## Discussion

The HIV-1 PPTs resist RNase H cleavage, function as RNA primers for plus-strand synthesis, and are displaced by RT to form the linear proviral dsDNA^2,6,11^ (Fig. 1). We report the first cryo-EM structures of HIV-1 RT bound to substrates containing either a PPT_DNA_ or PPT_RNA_ displacement strand plus incoming dATP. These structures reveal key features of the PPT displacement mechanism by HIV-1 RT. The displacement duplex exhibits an upward twist concomitant with a flip of the template nucleotide at the SD site, relocating the displacement strand more than 30 Å from the primer 3′-end (Fig. 3f). This dramatic repositioning underscores the mechanical requirements involved in displacement of the downstream PPT strand.

We propose that strand displacement depends on a “hydrophobic platform” created by cooperative interactions of W24 and F61 with template nucleotides T_0_, T_1_, and displacement nucleotide D_1_ at the SD site. Consistent with prior studies^45,64,65^, our data show F61 interacting critically with the template to coordinate translocation. Specifically, F61A impairs primer extension on all substrates, regardless of the presence of a displacement strand, while F61L rescues polymerization across substrates (Fig. 4b-d), indicating that the hydrophobic contacts at position 61 are essential for primer strand elongation.

F61 is also required for the SD activity, which depends on the ongoing DNA synthesis^22,24^. F61A reduces SD of both PPT_DNA_ and PPT_RNA_ (Fig. 4c-d), likely by disrupting the F61-T_1_ interaction needed to “unzip” T_1_ from D_1_ and translocate the template into the active site. F61L only partially rescues SD because leucine’s hydrophobic interaction cannot fully stabilize T_1_ in a “pulled-apart” conformation and guide it into the active site. However, F61L fully restores infectivity and ERT activity due to the extended timescales in these assays, aligning with an earlier finding^45^. Notably, 3DVAs reveal that the movement of F61 among various structural intermediates is concomitant with synchronized repositioning of the template strand T_1_ (Supplementary Videos 1, 2).

W24, on the other side of the platform, stacks specifically with the bases of T_1_ and D_1_. Structural comparisons with previously determined RT/T-P/nucleotide ternary complexes show that W24-T_1_ interactions vary due to the flexibility of the single-stranded template overhang (Supplementary Fig. 10a-c). However, the presence of the displacement strand at the SD site anchors the template through the W24-T_1_ interaction, thus stabilizing the complex in a pre-displacement state (Fig. 6a-b, Supplementary Fig. 10b-c). While W24 is proximal to T_1_ in some structures bearing single-stranded template overhang, biochemical characterization of W24A shows that W24 has no role in standard polymerization (Supplementary Fig. 10b-c, Fig. 4b). However, when encountering a downstream PPT strand, primer-elongation by W24A stalls upon reaching the displacement sites of the PPT_DNA_ and PPT_RNA_ strand (Fig. 4c-d). Furthermore, Viruses carrying W24A mutation are non-infectious, and ERT assays show defective late-stage RT products (Fig. 5b-c), consistent with impaired displacement. W24F fully rescues all activities confirming the need for an aromatic side chain at this position (Fig. 4c-d, Fig. 5b-c).

**Figure 6:**
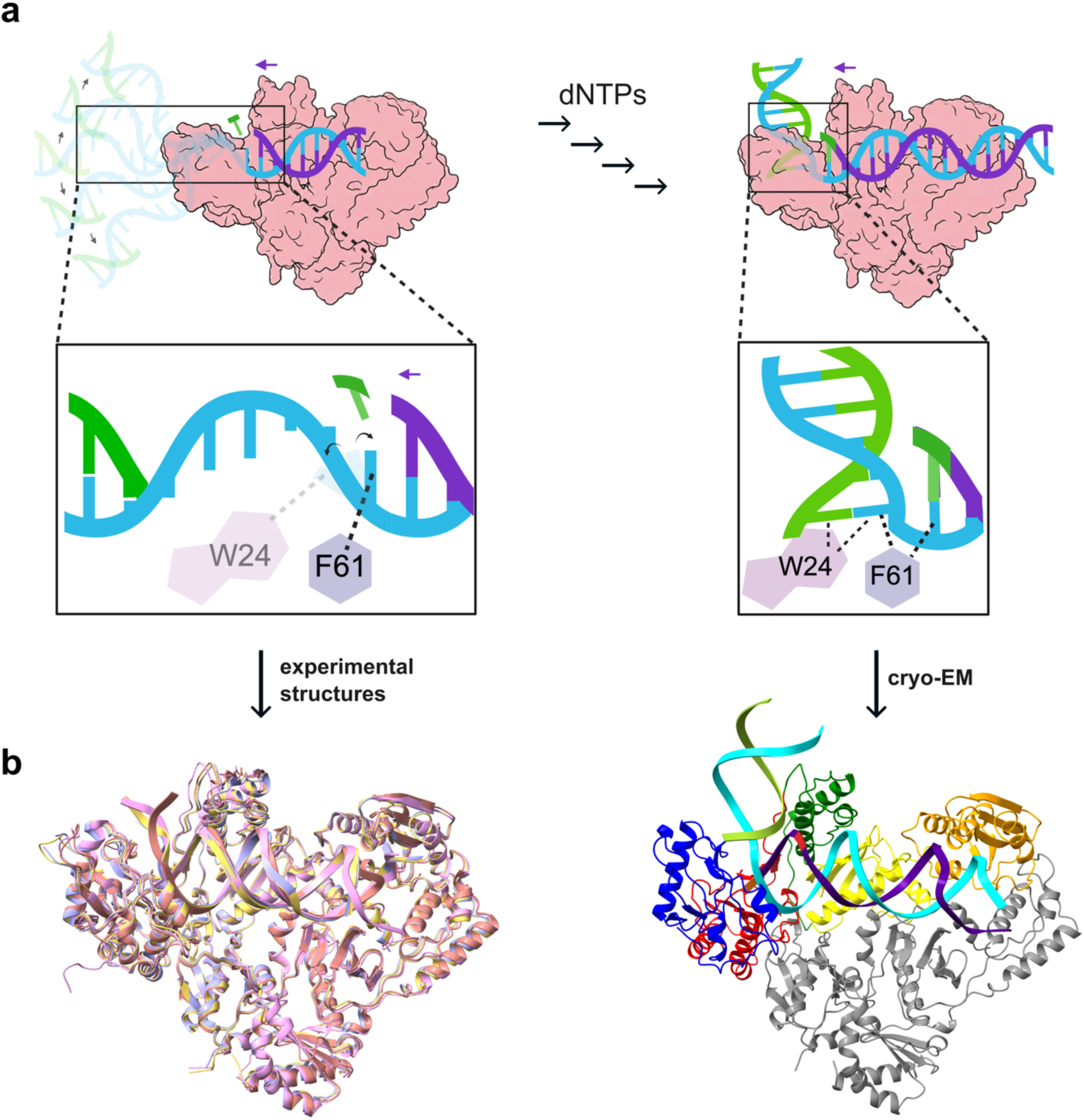
Strand displacement mechanism by HIV-1 RT. **a** Schematics of the polymerization and strand displacement mechanisms with zoomed-in views of the W24 and F61 coordination. **Left:** Structural comparisons show that the single-stranded portion of the template is highly flexible, and F61 interacts with the first nucleotide of the template overhang (T1). W24-T1 interactions vary depending on the position of the overhang. **Right:** The RT/T-P-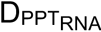/dATP complex shows RT in a pre-displacement state: the presence of the displacement strand locks the T-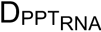 duplex in an upward-twisted orientation, coordinated by W24 and F61. **b** Experimental structures supporting the mechanisms proposed in **a**. **Left:** Structural superposition of the RT/T-P-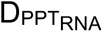/dATP complex with X-ray crystal structures of the RT/T-P/EFdA-TP (PDB: 5J2M; yellow), RT/T-P/dATP (PDB: 5TXL; pink), and RT/T-P/d4T-TP (PDB: 6WPF; light blue) complexes, as well as the cryo-EM structure of the RT/T-P/dATP complex (PDB: 8VB7; pastel red). Comparison details shown in **Supplementary Fig. 10**. **Right:** Cryo-EM structure of the RT/T-P-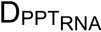/dATP complex. The RT p66 subunit is colored by subdomain: fingers (blue), palm (red), thumb (green), connection (yellow), and RNase H (orange). The p51 subunit is in gray. The primer strand is purple, the template strand is cyan, and the PPT-RNA displacement strand is light green. Schematics created in BioRender. Wen, X. (2026) https://BioRender.com/ydovuyh.

Double-mutant analysis suggests synergistic action by W24 and F61. The W24A/F61A double mutant abolishes both polymerization and SD in biochemical and viral assays (Figs. 4-5). Single-rescue mutants (W24A/F61L and W24F/F61A) fail to restore WT activities (Fig. 4b-d, Fig. 5b, d), indicating that W24 and F61 operate as a unified hydrophobic unit rather than independent residues. Only the double-rescue W24F/F61L restores full activity across assays (Fig. 4b-d, Fig. 5b, d), showing that the complete platform is required to coordinate template and displacement strands and minimize stalling during SD. Based on these findings, we propose a structural mechanism for HIV-1 RT strand displacement (Fig. 6a). During DNA-dependent DNA polymerization, the upstream single strand template remains flexible. At this stage, only the F61-T_1_ interaction is essential, whereas the W24-T_1_ interaction is not (Fig. 6a, left panel). As RT reaches the displacement strand, the downstream duplex is locked into an upward-twisted orientation, coordinated by W24 and F61 interacting with the displacement and template nucleotides (Fig. 6a, right panel). This more constrained pre-displacement state is exactly what is captured by our cryo-EM experiments. Furthermore, 3DVAs reveal that F61 shifts with the template strand across the captured conformations, whereas W24 remains relatively fixed at the SD site. This supports F61 is coordinating template translocation while W24 stabilizes the displacement strand (Supplementary Videos 1-2).

Additionally, R78 stabilizes the template strand by interacting with the phosphate backbone between T_1_ and T_0_. Biochemically, R78A specifically impairs SD on both PPT_DNA_ and PPT_RNA_ without affecting standard polymerization (Fig. 4b-d). In virions, however, R78A suppresses all RT products, and in ERT assays reduction starts from early stages, indicating broader defects in cDNA synthesis and reduced fitness (Fig. 5b-c). Although the R78K rescues biochemical SD, it fails to fully restore viral function (Fig. 4b-d, Supplementary Fig. 9c), highlighting arginine’s unique guanidinium group for bidentate phosphate interactions (versus lysine’s flexible primary amine side chain). Although the consensus map positions R78 approximately 3.7 Å from the template phosphate backbone, our 3DVAs show flexibility, with R78 contacting the template in certain conformations (Supplementary Videos 1-2). Thus, R78 likely acts as an electrostatic guide for T_1_, relieving torsional stress on the T_1_/T_0_ backbone and ensuring precise template trajectory during translocation and displacement.

PPT displacement faces a higher energetic barrier compared to standard DNA polymerization because the RNA/DNA hybrid adopts a rigid intermediate A/B-form conformation that is more stable than DNA/DNA B-form complexes^10,35,38,66^. This rigidity keeps the PPT duplex fully annealed in our structures, unlike the spontaneous unwinding seen in a human Pol γ strand displacement complex^59^. Displacing PPT_RNA_ therefore requires overcoming greater resistance to unzipping the hybrid^10,38^. Our biochemical assays show lower displacement efficiency for PPT_RNA_- compared to PPT_DNA_-containing substrates, reflecting these intrinsic differences (Fig. 4c-d, Supplementary Figs. 7-8). The different helical geometries were also captured in the cryo-EM structures at the downstream displacement site: while our PPT_DNA_ consensus map revealed a weak interaction between the displacement strand and T290 in the p66 thumb, the distinct coiling of the PPT_RNA_ shifted it away (Fig. 3d-e). We initially hypothesized that this differential T290 interaction contributed to the observed displacement kinetic differences. However, 3D classifications and 3DVAs revealed that the T290 interaction is transient in both substrates, appearing only in a particle subset rather than as a fixed state (Supplementary Fig. 5, Videos 1-2). The observation that T290 mutations had negligible effects on viral fitness and strand displacement efficiency for both substrates confirms that this contact is auxiliary rather than catalytic (Fig. 5b, Supplementary Fig. 7-8). Thus, the observed difference between PPT_DNA_ and PPT_RNA_ displacement strands is likely due to the intrinsic differences in DNA/DNA and RNA/DNA coiling rather than specific RT-substrate interactions. While the p66 thumb subdomain likely provides a broad channel to accommodate the downstream duplex, the primary mechanical determinants of strand displacement are localized to the fingers subdomain, where the specific residues (W24, F61, R78) at the SD site coordinate the energetic challenges of displacement duplex unzipping.

In summary, our structures and functional data define the mechanism of PPT strand displacement by HIV-1 RT, a critical step in late-stage reverse transcription required for linear dsDNA formation^1^. We propose that the conserved W24-F61 hydrophobic platform together with the R78 electrostatic guide forms a specialized displacement site beneath the template and displacement strands. This motif stabilizes displacement duplex geometry, facilitates unzipping, and directs template nucleotides into the polymerase active site for extension particularly important for the rigid PPT_RNA_/DNA hybrid. This mechanism enables full displacement of the cPPT to generate the central DNA flap essential for nuclear entry and pre-integration, and likely applies to the displacement of the identical-in-sequence 3′-PPT, completing the synthesis of the two LTRs necessary for viral integration and infectivity.

## Materials and Methods

### Plasmids and protein constructs

For structural studies, HIV-1 reverse transcriptase (RT) p66 and p51 subunits were expressed from pCDFDuet-1 and pRSFDuet-1 vectors, respectively. The p66 subunit contains the Q258C and C280S mutations, while the p51 subunit contains the C280S mutation; both constructs were cloned as previously described^67^. For biochemical assays, the pRT6H-PROT as previously described^68^.

### Oligonucleotides

For cryo-EM experiments, a modified primer strand (5′-ACAGTCCCTGTTCGGG*CGCC-3′), which contains a thioalkyl tether at the N2 position of G*, was used to enable covalent disulfide crosslinking to the RT p66 subunit at Q258C. The template strand was 5′-GCACTGTACCCCCCAATCCCCCCTTTTCTTTTTCGGCGCCCGAACAGGGACTGTG-3′. The polypurine tract (PPT) displacement strands used were either a DNA (5′-AAAAGAAAAGGGGGGATTGGGGGGTACAGTGC-3′) or an RNA-DNA strand (5′-rArArArArGrArArArArGrGrGrGrGrGATTGGGGGGTACAGTGC-3′). For biochemical assays, a 5′-Cy3-labeled primer (5′-Cy3-CAAATGGCAGTATTCATCC-3′) was annealed to the template strand 5′-GCACTGTACCCCCCAATCCCCCCTTTTCTTTTAAAATTGTGGATGAATACTGCCATTTG-3′. These assays used the same PPT displacement strands described above alongside a displacement-strand “trap” (5′-GCACTGTACCCCCCAATCCCCCCTTTTCTTTT-3′). The modified primer strand for crosslinking reaction was purchase from TriLink Biotechnologies, and all other oligonucleotides were obtained from Integrated DNA Technologies (IDT).

### Cell lines and viral vectors

HEK293T cells^69^ were purchased from Thermo Fisher Scientific, and TZM-GFP cells^61^ were generously provided by M. Pizzato (University of Trento). All cells were cultured and maintained in Dulbecco’s Modified Eagle Medium (DMEM; Mediatech) supplemented with 10% fetal bovine serum (FBS; VWR), 2 mM L-glutamine (ATCC), 1X non-essential amino acids (NEAA; Cytiva), and 100 U/mL penicillin/streptomycin (Gemini Bio). The viral envelope expression vector pVSV-G was obtained from Dr. Lung-Ji Chang via NIH BEI Resources (ARP-4693), and the Env-defective proviral clone pNL4-3ΔEnv-EGFP^70^ was obtain from Drs. Haili Zhang, Yan Zhou, and Robert Siliciano via NIH BEI Resources (ARP11100).

### Expression and Purification of HIV-1 RT

For cryo-EM experiments, HIV-1 RT p66 and p51 subunits were expressed and purified as previously described^52^. Briefly, pCDFDuet1 and pRSFDuet1 were overexpressed in BL21(DE3)-RIL *E. coli* in LB medium in the presence of streptomycin, kanamycin, and chloramphenicol to an OD_600_ of ∼0.7 at 37 °C, and expression was induced with 1 mM isopropyl β-d-1-thiogalactopyranoside (IPTG; VWR) for 3 h at 37 °C. Cells were harvested by centrifugation and lysed by sonication in the presence of lysozyme (Sigma Aldrich), phenylmethylsulfonyl fluoride (PMSF; Thermo Fisher Scientific), and trace amounts of DNase I (GoldBio) and RNase A (Roche). The clarified lysate was filtered and purified by nickel-affinity chromatography (Cytiva), followed by dialysis and heparin affinity chromatography (Cytiva) to separate free RT from nucleic-acid-bound RT. Fractions containing free RT were pooled, buffer-exchanged into storage buffer (10 mM Tris pH 8.0, 75 mM NaCl), concentrated to the desired concentration, aliquoted, and stored at −80 °C.

For biochemical experiments, RT was expressed and purified as described previously^60^. Briefly, pRT6H-PROT was overexpressed in JM109 *E. coli* (Invitrogen) in LB medium in the presence of chloramphenicol to an OD_600_ of ∼0.7 at 37 °C, and expression was induced with 1 mM IPTG for 3 h at 37 °C. Cells were harvested by centrifugation and lysed by sonication in the presence of lysozyme, PMSF, and trace amounts of DNase I and RNase A. The clarified lysate was filtered and incubated with 2 mL of pre-washed nickel-affinity resin (Sigma Aldrich) for 2 h at 4 °C. The resin was washed with buffer (50 mM sodium phosphate, pH 7.8, 300 mM NaCl) containing 20 mM imidazole, and the RT was eluted using the same buffer containing 250 mM imidazole. Purified RT was buffer-exchanged into storage buffer (50 mM Tris pH 8.0, 1 mM EDTA, 1 mM DTT, 25 mM NaCl, and 50% glycerol), concentrated to the desired concentration, aliquoted, and stored at −80 °C.

### Cryo-EM sample and grid preparations

To prepare the crosslinked RT-nucleic acid complex for cryo-EM experiments, the template, primer, and displacement strands (T-P-D_PPT_) were annealed at a 1:1:2 molar ratio (T:P:D) in annealing buffer (40 mM Tris-HCl, pH 8.0, 50 mM NaCl, 10 mM MgCl_2_) by heating to 95 °C for 5 min followed by slow cooling to room temperature for 40 min. The annealed T-P-D_PPT_ substrates (20 μM) were then incubated with crosslinkable HIV-1 RT (6.3 μM) in a buffer containing 25 mM Tris-HCl (pH 8.0), 100 mM NaCl, 10 mM MgCl_2_, and 0.1 mM 2′,3′-dideoxyguanosine-5′-triphosphate (ddGTP; GE Healthcare Life Sciences). Crosslinked complexes were purified using a nickel-heparin tandem chromatography method as described previously^52^. Fractions containing the crosslinked complex were pooled, buffer-exchanged into EM buffer (75 mM Tris pH 8.0, 150 mM NaCl), and concentrated to ∼10 mg/mL. Finally, the complex was incubated with 5 mM MgCl_2_, 240 μM dATP, and 0.2% β-octyl glucoside^71^ for 30 min at 4 C before plunge freezing.

For vitrification, the crosslinked complexes were applied on the Quantifoil 300-mesh R1.2/1.3 UltrAuFoil® grids. Grids were glow discharged in a Solarus system at −15 mA for 30 s; then, 3 μL of sample was applied and blotted for 3 s with a blot force of 0. Vitrification was performed by plunge-freezing into liquid ethane using a Vitrobot Mark IV with the chamber set to 22 °C and 100% humidity.

### Cryo-EM data collection and processing

Cryo-EM data were collected on a 300 kV Titan Krios G3i TEM (Thermo Scientific) equipped with a Ceta 16M CMOS camera and a Gatan K3 direct electron detector operated in super-resolution mode. Automated data acquisition was performed using EPU (Thermo Scientific). 50-frame movies were recorded at a nominal magnification of 105,000X with a binning of 2, yielding a physical pixel size of 0.825 Å. The total electron dose was 50 e^-^/Å^2^, and data were collected over a defocus range of −0.8 to −2.0 μm in 0.1 μm increments.

Datasets were processed in RELION^72^ and CryoSPARC^73^. The detailed processing workflow is shown in Supplementary Figs. 1 and 3. Briefly, micrographs were motion-corrected in RELION using MotionCor2^74^ and then transferred to CryoSPARC for particle picking, 2D classifications, 3D reconstruction and refinements, and CTF corrections. Particle stacks were transferred back into RELION for Bayesian polishing^75^, further CTF corrections, and final 3D refinements.

### Model building and refinement

The structure of RT/T-P/EFdA-TP^52^ (PDB ID 5J2M) was used as the initial model for building the RT/T-P-D_PPT-DNA_/dATP complex. The additional displacement duplex DNA was built in UCSF ChimeraX^76,77^. Models were iteratively improved by manual rebuilding in Coot v0.9.8.95^78^ followed by real-space refinement in PHENIX v1.21.2^79^. The resulting model was then used as the starting model for the other PPT_DNA_ complexes, as well as the initial RT/T-P-D_PPT-<u>RNA</u>_/dATP complex. The RNA/DNA displacement duplex was built in UCSF ChimeraX. The final refined RT/T-P-D_PPT-<u>RNA</u>_/dATP complex was used as the initial model for the other PPT_RNA_ complexes. All models were iteratively improved by manual rebuilding in Coot followed by real-space refinement in PHENIX, as described above. Additional data collection and processing parameters are listed in Supplementary Table 1.

### Site-directed mutagenesis

Site-directed mutagenesis of the HIV-1 RT enzyme was performed using the pRT6H-PROT backbone and the Q5^®^ Site-Directed Mutagenesis Kit (New England Biolabs, NEB) according to the manufacturer’s instructions. Primer sequences for each RT mutant are listed in Supplementary Table 2. Following the PCR reaction and KLD treatment, products were transformed into JM109 *E. coli* and recovered in SOC medium for 1 h at 37 °C.

Mutations in the RT gene were also introduced into pNL4-3ΔEnv using PCR and the NEBuilder® HiFi DNA Assembly Master Mix (NEB) according to the manufacturer’s instructions. The assembly vector backbone was generated by digesting pNL4-3ΔEnv with ApaI and SalI restriction enzymes. Primer sequences for each RT mutant and restriction sites used are provided in Supplementary Table 3. PCR products for each mutant were resolved on 0.8% agarose gels, and the appropriate DNA fragments were gel-purified using the QIAquick Gel Extraction Kit (QIAGEN). Assembled products were treated with KLD mix, followed by transformation into DH5α cells (NEB).

For both mutagenesis strategies, transformed cells were plated on LB agar containing chloramphenicol for overnight incubation at 37 °C. Individual colonies were then cultured in LB medium with chloramphenicol, and plasmid DNA was isolated using the QIAprep Spin Miniprep Kit (QIAGEN) according to the manufacturer’s protocol. All mutations were confirmed by Sanger sequencing (Genewiz/Azenta Life Sciences).

### Biochemical primer-extension and strand-displacement assays

Wild-type (WT) and mutant HIV-1 RT (encoded by pRT6H-PROT) were expressed and purified as described above and used for the biochemical primer-extension assays. Expression from the pRT6H-PROT construct involves the simultaneous expression of RT_p66_ and the viral protease, which cleaves ∼ 50% of the RT_p66_ into RT ^68^. Heterodimeric RT (p66/p51) assembles in solution and is subsequently purified. Consequently, all produced mutants contain their corresponding mutations on both the p66 and p51 subunits. Oligonucleotide sequences used are also listed above. Nucleic acid substrates lacking or containing a downstream displacement strand were prepared by annealing the template and primer at a 1:2 molar ratio (T:P) or the template, primer, and displacement strand at a 1:2:3 molar ratio (T:P:D), respectively, in annealing buffer.

Primer extension reactions were performed similarly as described previously^60^. Briefly, 20 nM annealed substrate was incubated with HIV-1 RT (WT or mutant) in RT buffer containing 50 mM Tris (pH 7.8) and 50 mM NaCl, supplemented with 80 nM displacement-strand “trap” (sequence listed above). Reactions with the T-P and T-P-D_PPT-DNA_ substrates used 40 nM RT, whereas reactions with the T-P-D_PPT-RNA_ substrate used 80 nM RT. Reactions were initiated by the addition of 6 mM MgCl2 and 15 μM dNTPs to a final volume of 20 μL. Reactions containing the T-P and T-P-D_PPT-DNA_ substrates were quenched after 15 and 60 min, whereas reactions containing the T-P-D_PPT-RNA_ substrate were quenched after 30 and 90 min, by adding 20 μL of 100% formamide containing trace of bromophenol blue.

Reaction products were resolved on 15% polyacrylamide/7 M urea gels and scanned using a Typhoon FLA 9500 (GE Healthcare). Band intensities were quantified with AzureSpot Pro (Azure Biosystems). Bands of interest were classified as no extension (position 0), pre-displacement (position 8), early-displacement (positions 9-13), mid-PPT displacement (positions 16-18), and full extension (position 40) (detailed in Fig. 4). The intensity of each bands-of-interest was normalized to the total intensity of the corresponding lane. Normalized intensities from the five listed positions were summed to represent 100% intensity. The percentage of each product was then calculated relative to this sum and plotted using GraphPad Prism 10. Statistical analyses for each position of interest were performed using a one-way ANOVA with Dunnett’s multiple comparisons test, comparing each mutant to the WT control on the corresponding gel.

### Pseudotyped VSV-G-HIV-1 virion production

The pVSV-G and pNL4-3ΔEnv (encoding either WT or mutant RT) plasmids were co-transfected into HEK293T cells at a 1:5 plasmid ratio using the X-tremeGENE HP transfection reagent (Roche) in Opti-MEM (Gibco). Viral supernatants were harvested 48 h post-transfection and filtered through a 0.45 μm polyethersulfone (PES) membrane (Millipore). Virions were concentrated by centrifugation at 14,000 rpm for 2.5 h, resuspended in PBS, aliquoted, and stored at −80 °C.

### Capsid (CA) p24 enzyme-linked immunosorbent assay (ELISA)

Viral yield from each transfection was quantified by measuring p24 levels using an in-house ELISA as previously described^80^. High-binding 96-well plates (Costar 3922, Corning) were coated with anti-p24 antibody (Abcam, ab9071) and blocked with 3% bovine serum albumin (BSA). Each batch of the VSV-G-pseudotyped HIV-1 virions was lysed and diluted in a buffer containing 0.5% BSA and 0.1% Triton X-100, added to the plates, and incubated for 1 h. A standard curve was generated in parallel using known concentrations of in-house purified p24^81^. Plates were washed with PBS-T, incubated with primary antibody (HIV-Ig; HIV Reagent Program, ARP-3957), washed, and incubated with an HRP-conjugated anti-human secondary antibody (AB_2337532; Jackson ImmunoResearch). After another PBS-T wash, chemiluminescent substrate (Thermo Fisher Scientific) was added, and the signal was measured on a GloMax^®^ microplate reader (Promega). p24 concentrations for each viral batch were then calculated from the standard curve.

### p24 normalized infectivity

Equal amounts of virions (encoding either WT or mutant RT), normalized to p24 levels, were used to infect TZM-GFP cells. Infection efficiency was quantified by counting GFP-positive cells 48 h post-infection (hpi) using the Cytation 5 Cell Imaging Multimode Reader with Gen5 software (BioTek). Data were analyzed using GraphPad Prism 10.

### Endogenous reverse transcription (ERT) assay

The same batches of p24-normalized virions (encoding either WT or mutant RT) used for the infectivity assays were also used for the ERT assay. ERT was performed as previously described^62,63,80^. Briefly, equal amounts of virions (based on p24 levels) were incubated overnight at 37 °C in the presence of melittin (2.5 μg/mL), BSA (0.5 mg/mL), IP_6_ (80 μM), and dNTPs (40 μM) in a reaction buffer containing 10 mM Tris-HCl (pH 7.8), 75 mM NaCl, and 2 mM MgCl_2_. Products corresponding to defined steps of HIV-1 reverse transcription were quantified by SYBR Green qPCR (Applied Biosystems) performed on the QuantStudio 3 Real-Time PCR System (Thermo Fisher Scientific). Serial dilutions of the pNL4-3ΔEnv plasmid DNA were used to generate standard curves for absolute quantification. For each mutant, the number of DNA copies calculated for each reverse transcription product was normalized to the corresponding product of the WT control. Data were analyzed using GraphPad Prism 10. Primer sets used to detect each reverse transcription products are detailed in Fig. 1 and as follows: minus-strand strong stop [(−)ssSTOP]: 5’-GCCTCAATAAAGCTTGCCTTGA-3’ and 5’-TGACTAAAAGGGTCTGAGGGATCT-3’; first-strand transfer (FST): 5’-GAGCCCTCAGATGCTGCATAT-3’ and 5’-CCACACTGACTAAAAGGGTCTGAG-3’; full-length minus-strand (FLM): 5’-CTAGAACGATTCGCAGTTAATCCT-3’ and 5’-CTAT CCTTTGAT GCACACAATAGAG −3’; second strand transfer (SST): 5’-TGTGTGCCCGTCTGTTGTGT-3’ and 5’-GAGTCCTGCGTCGAGAGATC-3’.

## Supporting information

Descriptions for Supplementary Videos

## Acknowledgement

This research was supported in part by the National Institutes of Health (R37 AI076119 to S.G.S.; F31 AI179424 to X.W.; F31 AI174951 to W.M.M.; W.M.M. was supported in part by T32 GM135060. Cryogenic Electron Microscopy (cryo-EM) was carried out by the Laboratory for BioMolecular Structure (LBMS) at the Brookhaven National Laboratory (BNL). The LBMS is supported by the DOE Office of Biological and Environmental Research (KP1607011). Cryo-EM condition optimization experiments were carried out by Emory University Robert P. Apkarian Integrated Electron Microscopy Core Facility (IEMC; RRID: SCR_023537), which is subsidized by the School of Medicine and Emory College of Arts and Sciences. Additional support was provided by the Georgia Clinical & Translational Science Alliance of the National Institutes of Health under award number UL1TR000454. We acknowledge Emory IEMC personnel Drs. Ricardo C. Guerrero-Ferreira and Srihari Nagendra Ravi K. Koripella for their technical support. The content is solely the responsibility of the authors and does not necessarily represent the official views of the National Institutes of Health. Additionally, S.G.S. acknowledges funding from the Nahmias-Schinazi Distinguished Chair in Research.

## Contributions

X.W. and S.G.S. designed the project and wrote the manuscript. X.W. purified and assembled the complexes, prepared cryo-EM grids for cryo-EM collection. W.M.M. established the initial computational infrastructure and server architecture used for preliminary cryo-EM data processing. X.W. and R.A.D. performed cryo-EM data processing. With guidance from R.A.D., K.A.K., and S.G.S., X.W. performed all model buildings and refinements. X.W. and R.L. performed mutagenesis and cloning for all RT-mutant constructs. X.W. purified enzymes for biochemical studies and conducted *in vitro* primer-extension and strand-displacement assays; X.W. and SD.M. performed and analyzed the associated PAGE experiments. X.W. and R.L. generated pseudotyped HIV-1 virions, performed and analyzed p24 ELISA and viral infectivity assays. X.W. performed and analyzed endogenous reverse transcription assays. All authors contributed to manuscript editing.

---

**Supplementary Fig. 1.**
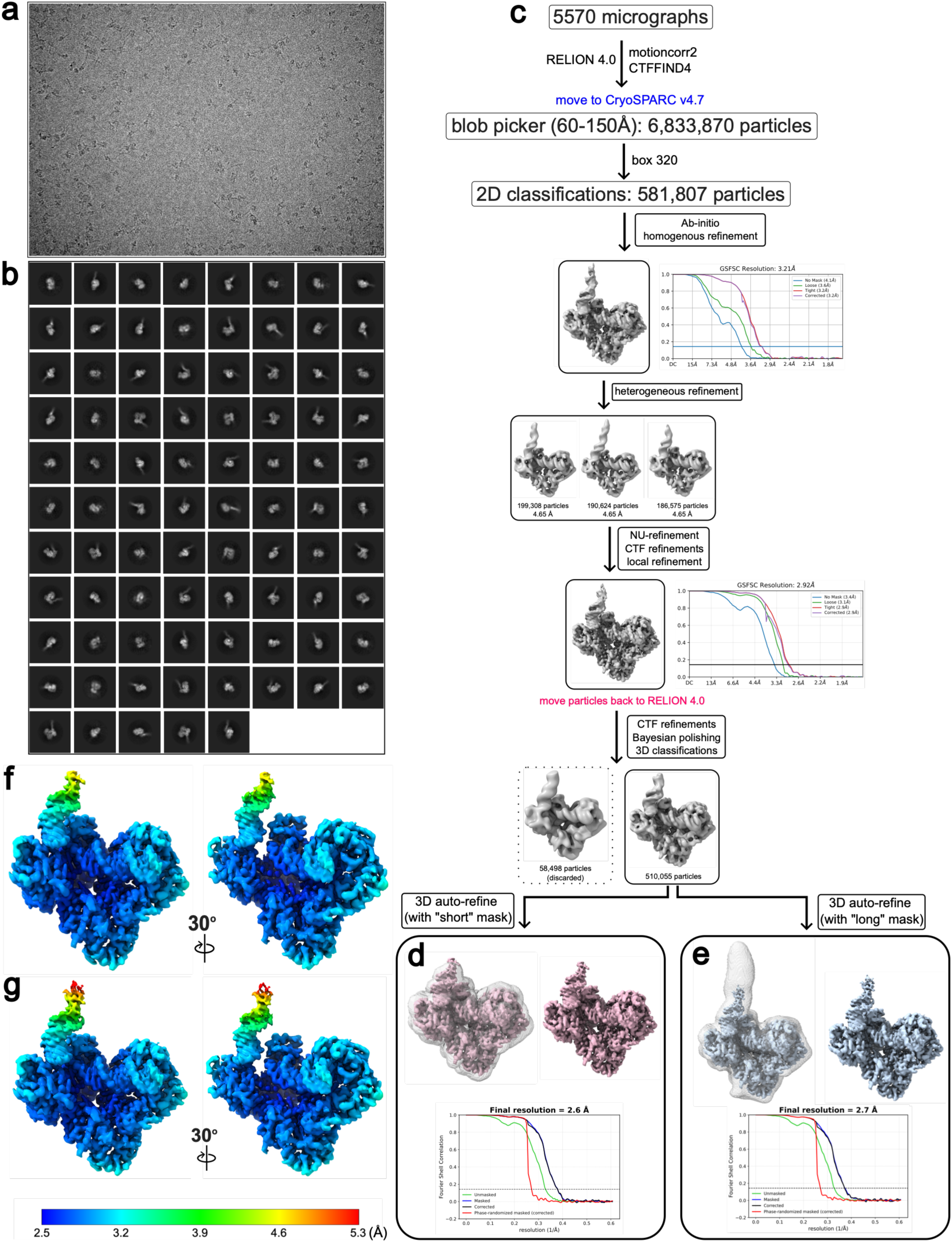
Workflow of cryo-EM processing of the RT/T-P-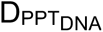/dATP complex. **a** Representative cryo-EM micrograph. **b** 2D classes. **c** Cryo-EM data processing workflow performed in CryoSPARC v4.7 and RELION 4.0. The number of particles selected at each stage is indicated. **d** Final 3D auto-refined map (unsharpened) contoured at 6.7σ and corresponding Gold-standard Fourier Shell Correlation (FSC) curve using a “short” mask. The final global resolution is 2.6 Å at the FSC = 0.143 criterion. **e** Final 3D auto-refined map (unsharpened) contoured at 6.0σ and corresponding Gold-standard FSC curve using a “long” mask. The final global resolution is 2.7 Å at the FSC = 0.143 criterion. **f-g** Local resolution estimates for the maps shown in **d** and **e**, respectively. The color bar indicates resolution range values in Å.

**Supplementary Fig. 2.**
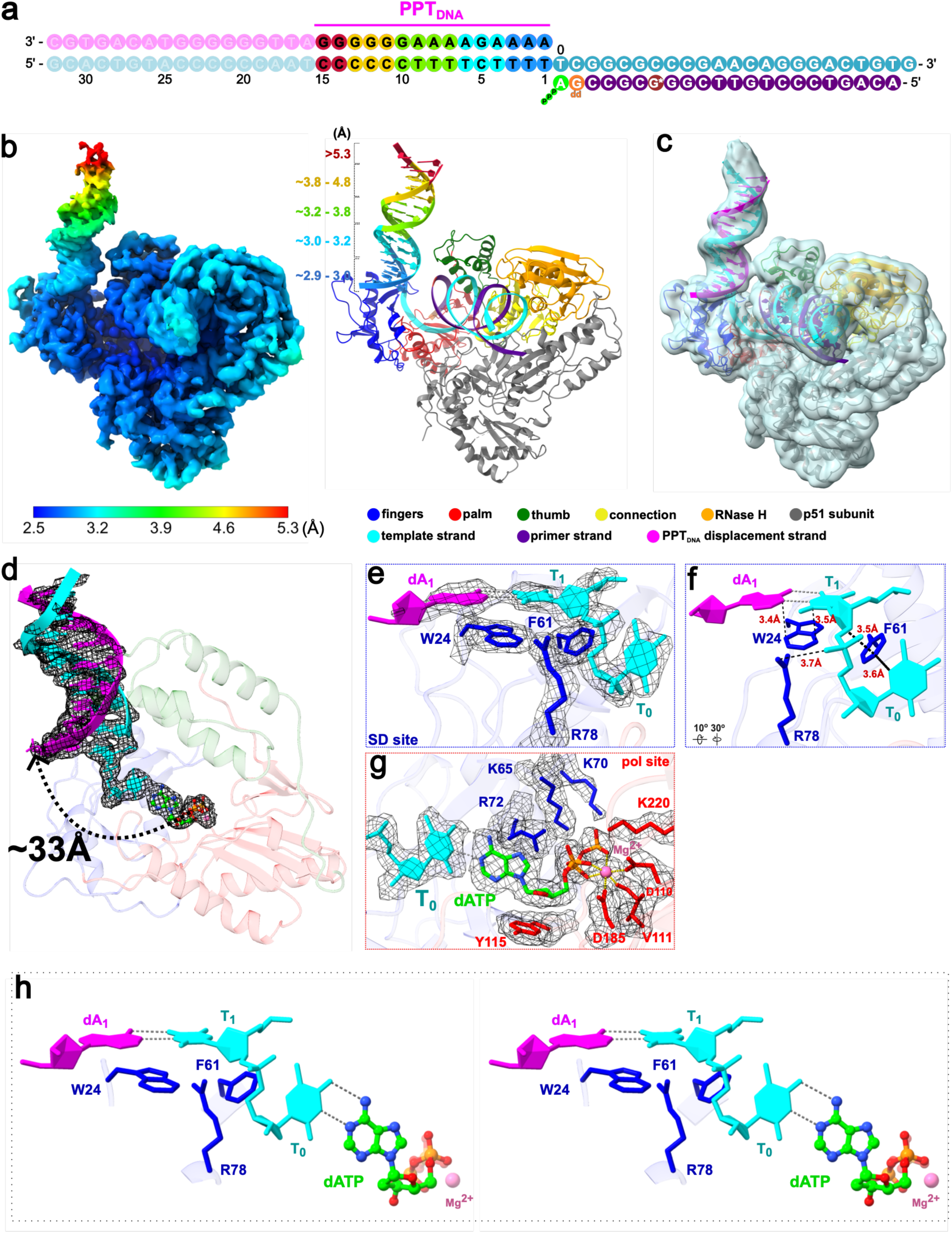
Cryo-EM structure of HIV-1 RT in complex with PPT_DNA_ displacement substrate and incoming dATP (RT/T-P-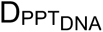/dATP). **a** Schematic of the nucleic acid substrate. The primer strand (purple) contains modifications and coloring as described in Fig. 2. The multicolor gradient on the PPT_DNA_ portion of the displacement duplex corresponds to the local resolution range observed in the EM density, as indicated by the color bar in **b**. Faded regions indicate segments not resolved. **b** Cryo-EM density map (**left**) and atomic model (**right**) of the RT/T-P-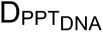/dATP complex with the extended 15-bp displacement duplex. The density map was refined using the “long” mask (**Supplementary Fig. 1e**) and is contoured at 6.0σ. Local resolution colors of the displacement duplex are mapped onto the atomic model as indicated in **a**. The color bar indicates resolution range values in Å. **c** Gaussian low-pass filtered map (σ=1.3 Å) contoured at 5.4σ (light blue), was used to guide the modeling of the extended displacement duplex. **d** Close-up view of the downstream displacement duplex and incoming dATP (corresponding to Fig. 2f). The cryo-EM density (mesh) from the 3D-refined map (unsharpened) is contoured at 8.1σ. The approximate distance (∼33 Å) between the α-phosphate of the incoming dATP and the O5’ atom of the first displacement strand nucleotide (dA1) is indicated. **e** Cryo-EM density (mesh) from the postprocessed map, overlaid on the atomic model of the strand displacement (SD) site shown in Fig. 2g. The map is contoured at 10.9σ. **f** View of the SD site rotated relative to **e**. **g** Cryo-EM density (mesh) from the postprocessed map, overlaid on the atomic model of the polymerase active site shown in Fig. 2h. The map is contoured at 7.1σ. **h** Stereo view (wall-eyed) of the SD site residues and the incoming dATP. Distances are shown as black dash lines, and hydrogen bonds are shown as gray dash lines.

**Supplementary Fig. 3.**
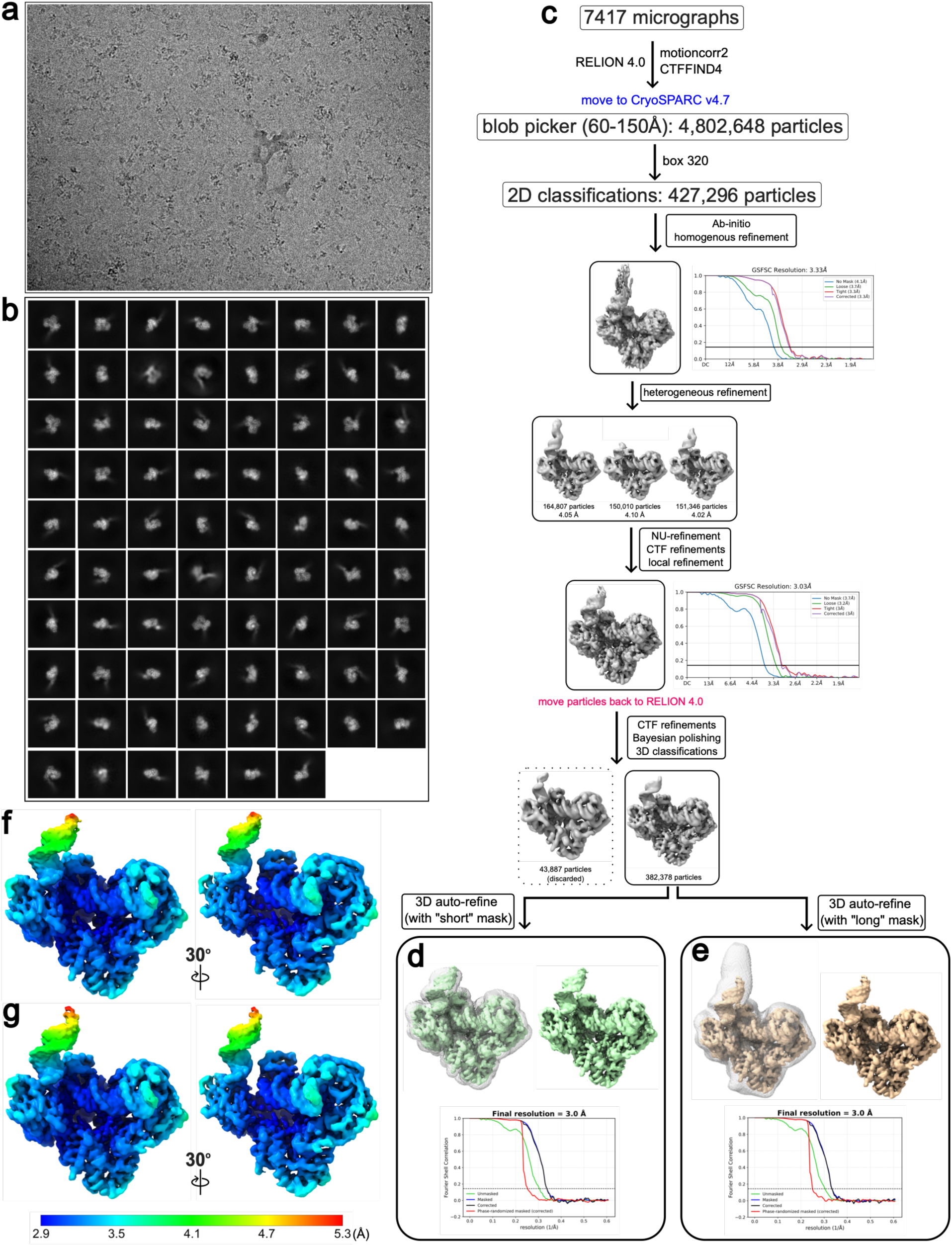
Workflow of cryo-EM processing of the RT/T-P-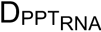/dATP complex. **a** Representative cryo-EM micrograph. **b** 2D classes. **c** Cryo-EM data processing workflow performed in CryoSPARC v4.7 and RELION 4.0. The number of particles selected at each stage is indicated. **d** Final 3D auto-refined map (unsharpened) contoured at 7.4σ and corresponding Gold-standard Fourier Shell Correlation (FSC) curve using a “short” mask. The final global resolution is 3 Å at the FSC = 0.143 criterion. **e** Final 3D auto-refined map (unsharpened) contoured at 6.6σ and corresponding Gold-standard FSC curve using a “long” mask. The final global resolution is 3 Å at the FSC = 0.143 criterion. **f-g** Local resolution estimates for the maps shown in **d** and **e**, respectively. The color bar indicates resolution range values in Å.

**Supplementary Fig. 4.**
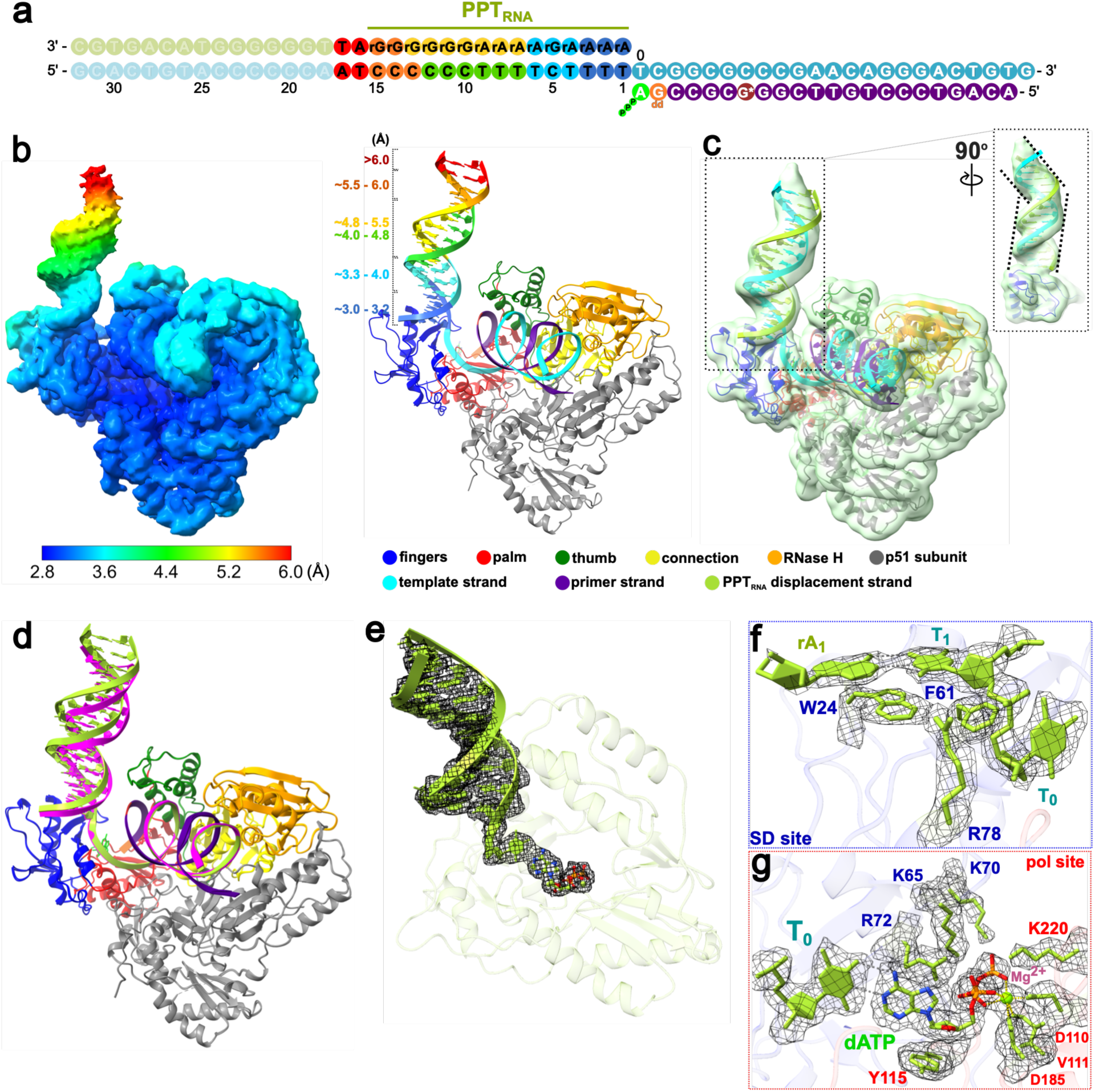
Cryo-EM structure of HIV-1 RT in complex with PPT_RNA_ displacement substrate and incoming dATP (RT/T-P-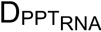/dATP). **a** Schematic of the nucleic acid substrate. The primer strand (purple) contains modifications and coloring as described in Fig. 2. The multicolor gradient on the PPT_DNA_ portion of the displacement duplex corresponds to the local resolution range observed in the EM density, as indicated by the color bar in **b**. Faded regions indicate segments not resolved. **b** Cryo-EM density map (**left**) and atomic model (**right**) of the RT/T-P-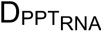/dATP complex with the extended 17-bp displacement duplex. The density map was refined using the “long” mask (**Supplementary Fig. 2e**) and is contoured at 5.2σ. Local resolution colors of the displacement duplex are mapped onto the atomic model as indicated in **a**. The color bar indicates resolution range values in Å. **c** Gaussian low-pass filtered map (σ=1.3 Å; light green), contoured at 5.4σ, used to guide modeling of the extended 17-bp PPT_RNA_ displacement duplex. Right: a 90° rotated view of the PPT_RNA_ displacement duplex, with black dashed lines indicating its helical axis directions. An apparent bend occurs within the region of consecutive rG of the PPT_RNA_ sequence. **d** Structural overlay of the extended (15-bp) RT/T-P-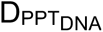/dATP and extended (17-bp) RT/T-P-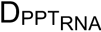/dATP complexes. Enzyme and primer-strand are colored as described previously. The template and displacement strands are colored magenta for the PPT_DNA_ model and yellow-green for the PPT_RNA_ model. **e** Close-up view of the downstream displacement duplex and incoming dATP (corresponding to the view in Fig. 3f). The cryo-EM density (mesh) from the 3D-refined map (unsharpened) is contoured at 7.4σ. **f** Cryo-EM density (mesh) from the post-processed map, overlaid on the atomic model of the strand displacement (SD) site shown in Fig. 3g. The map is contoured at 7.2σ. **g** Cryo-EM density (mesh) from the post-processed map, overlaid on the atomic model of the polymerase (pol) active site shown in Fig. 3h. The map is contoured at 5σ. Dashed gray lines indicate hydrogen bonds.

**Supplementary Fig. 5.**
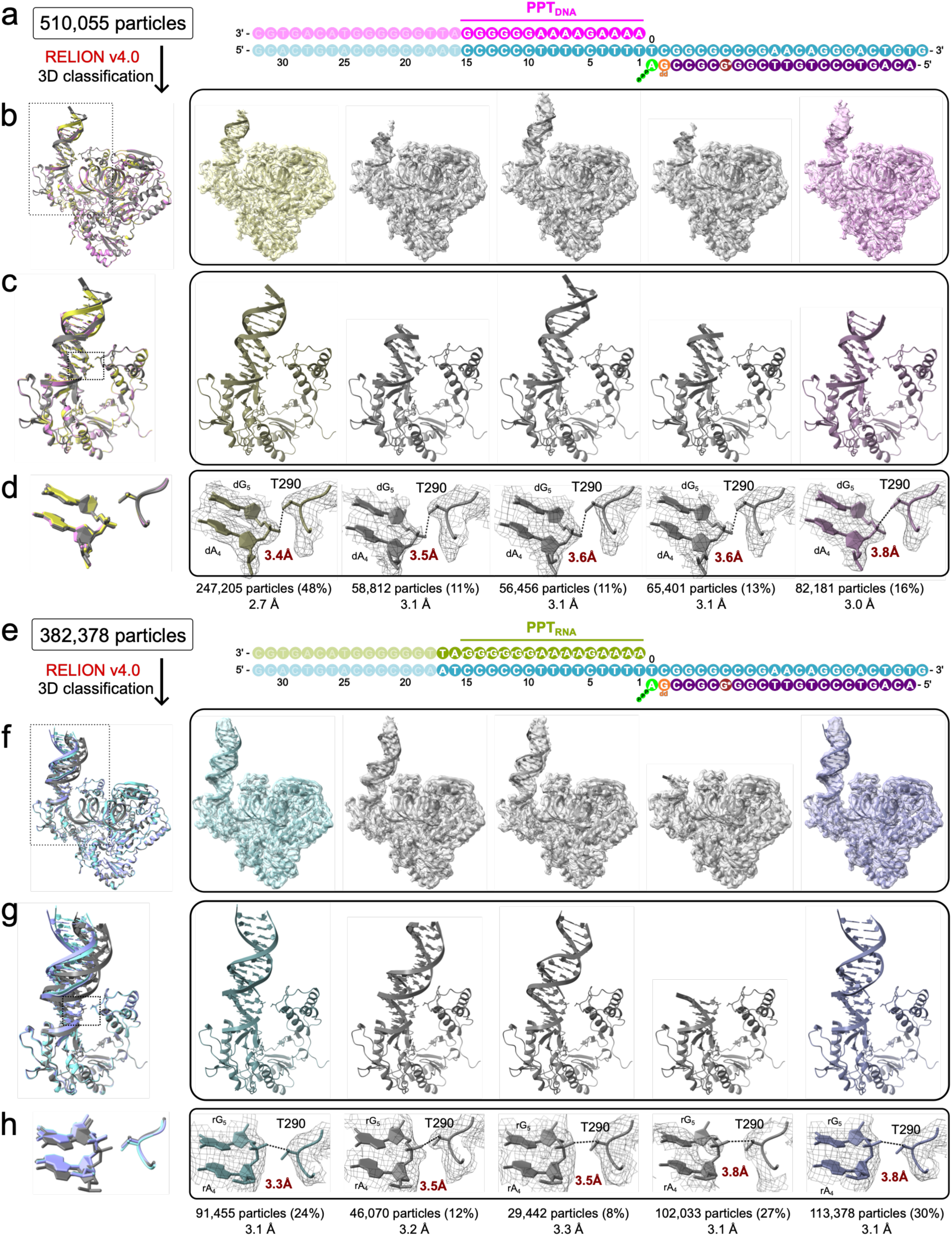
3D classifications of the final particle stacks for the RT/T-P-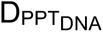/dATP and RT/T-P-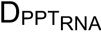/dATP complexes. **a-d** 3D classification (5 classes) of the final particle stack for the RT/T-P-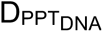/dATP complex. **e-h** 3D classification (5 classes) of the final particle stack for the RT/T-P-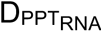/dATP complex. **a, e** Left: number of particles used for 3D classification. Right: schematic of the nucleic acid substrate. **b, f** Overlay of the atomic models for the five 3D classes. **Right panels**: Individual classes showing atomic models fitted into their corresponding unsharpened 3D-refined maps. All maps are shown at an absolute contour level of 0.0055. **c, g** Close-up view (boxed region in **b, f**) showing the overlay of the polymerase subdomain and the downstream displacement duplex. **Right panels**: Individual classes at this view. **d, h** Close-up view (boxed region in **c, g**) detailing the contact between T290 and the displacement strand. **Right panels**: Individual classes at this view, with the distance between T290 and the dG_5_ (**d**) or rG_5_ (**h**) phosphate indicated for each. The corresponding unsharpened 3D-refined density maps (mesh) are shown at an absolute contour level of 0.0055 (except 0.0035 for class 4 in **h**). The class exhibiting the closest T290 contact with the PPT_DNA_ (PDB 11XI, EMDB 76156) is colored yellow (**d**); the closest T290 contact with the PPT_RNA_ (PDB 11XM, EMDB 76160) is colored light blue (**h**). The class with the furthest T290-PPT_DNA_ contact (PDB 11XJ, EMDB 76157) is colored pink (**d**); the furthest T290-PPT_DNA_ contact (PDB 11XN, EMDB 76161) is colored light purple (**h**). All other classes with intermediate contacts are gray. The number of particles, percentage of the total stack, and final map resolution are indicated for each class.

**Supplementary Fig. 6.**
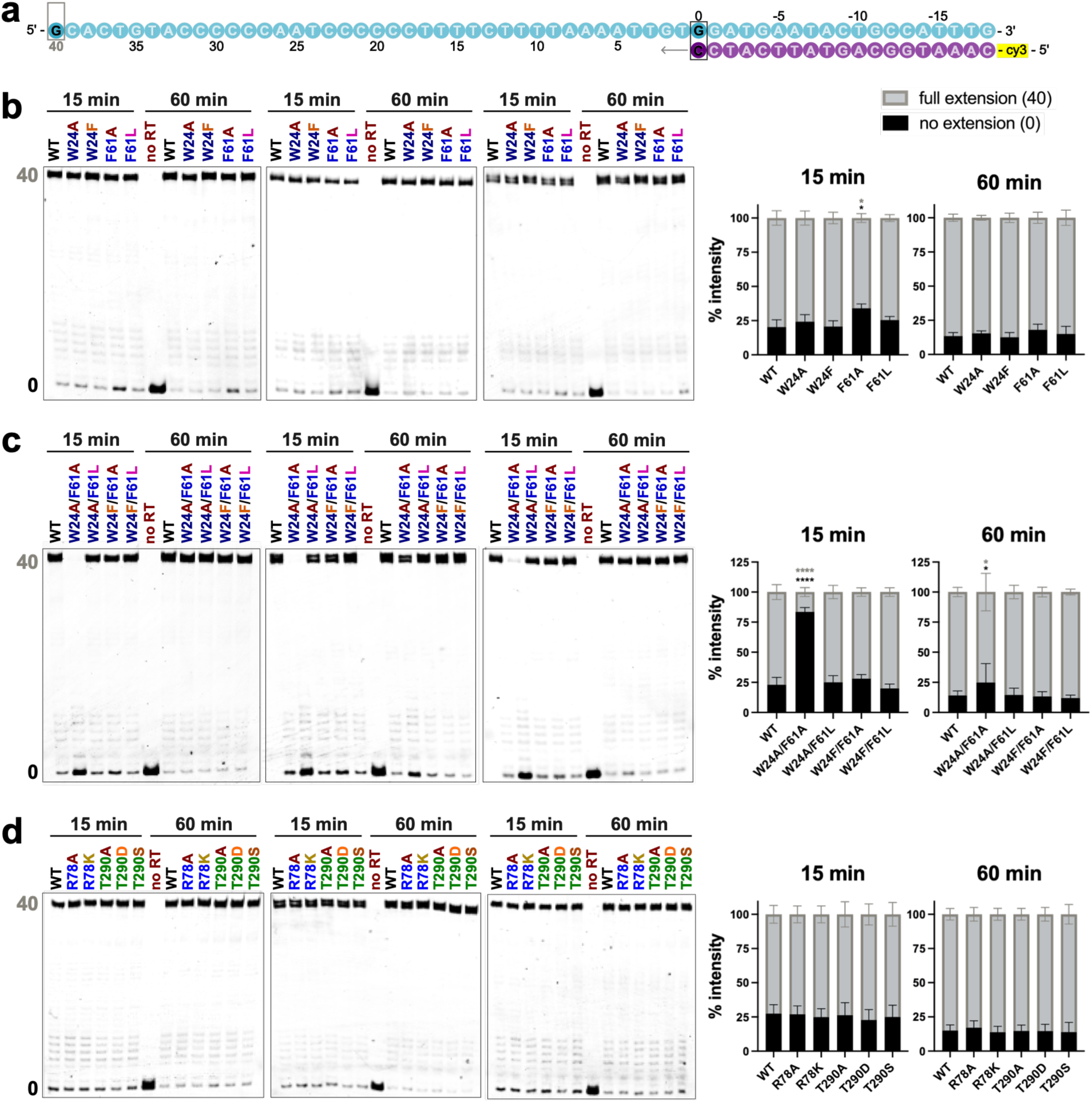
Triplicate primer-extension assays using the template-primer (T-P) only substrate. **a** Schematic of the T-P only nucleic acid substrate. The primer strand (purple) is labeled with a Cy3 fluorophore at the 5’-end, and the template strand is cyan. Boxed, darker-colored nucleotides represent strong primer-extension pause sites observed in the assays. **b-d** Triplicate PAGE gels and corresponding quantitative bar graphs for primer-extension assays performed with single W24 and F61 mutants (**b**), double W24 and F61 mutants (**c**), and single R78 and T290 mutants (**d**) after 15- and 60-min reactions. Gel bands represent the size of the Cy3-labeled primer. Quantified positions of interest include the un-extended primer at position 0 (black) and the fully extended primer at position 40 (gray). Statistical significance was determined using a one-way ANOVA with Dunnett’s multiple comparisons test, comparing each mutant to the WT control on the corresponding gels. P < 0.05; ****P < 0.0001; bars without asterisks are not significant (P > 0.05).

**Supplementary Fig. 7.**
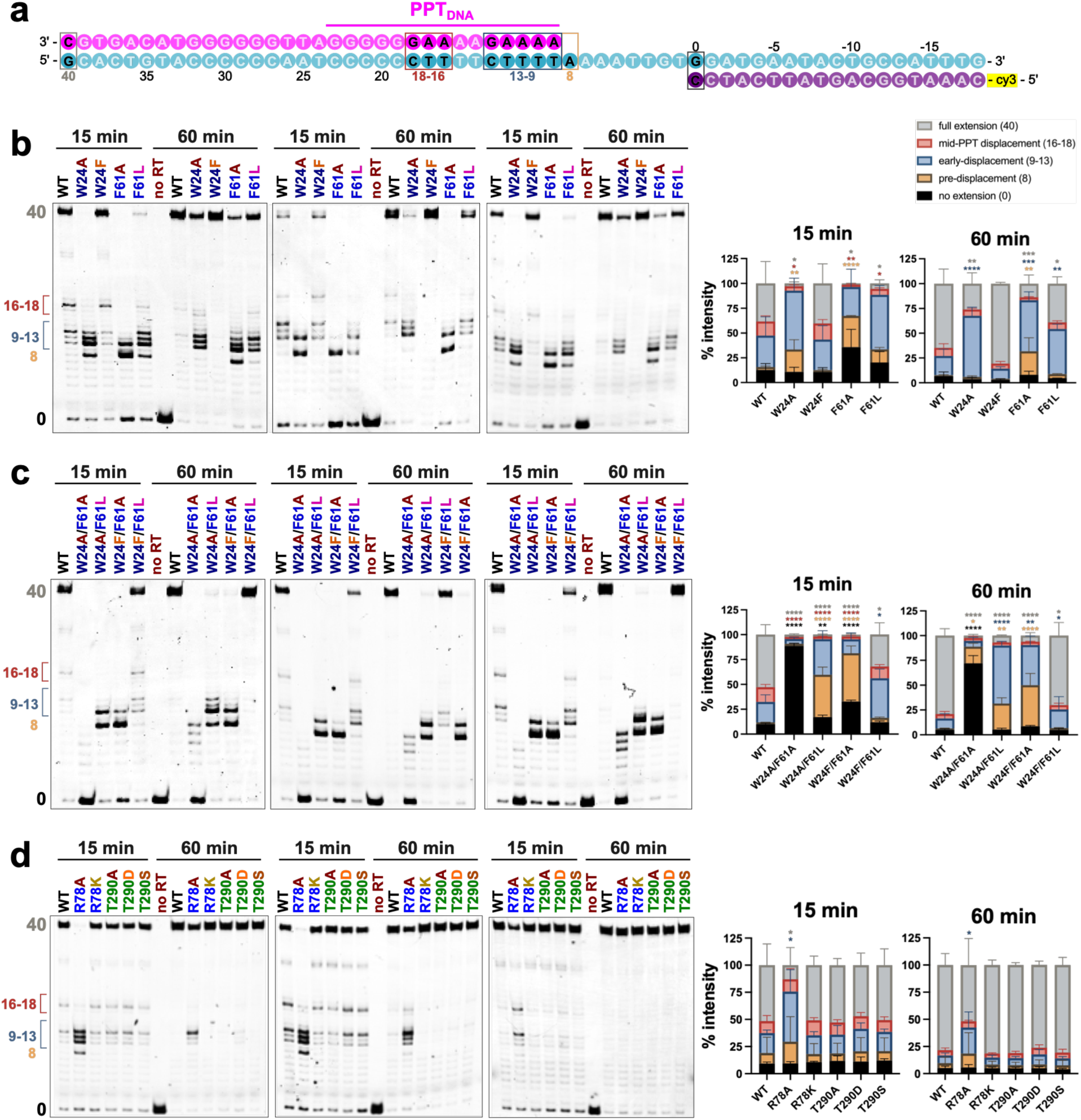
Triplicate primer-extension assays using the PPT_DNA_ displacement (T-P-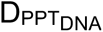<u>_)_</u> substrate. **a** Schematic of the T-P-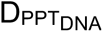 nucleic acid substrate. The primer strand (purple) is labeled with a Cy3 fluorophore at the 5’-end, the template strand is cyan, and PPT_DNA_ displacement strand color in magenta. Boxed, darker-colored nucleotides represent strong primer-extension pause sites observed in the assays. **b-d** Triplicate PAGE gels and corresponding quantitative bar graphs for primer-extension assays performed with single W24 and F61 mutants (**b**), double W24 and F61 mutants (**c**), and single R78 and T290 mutants (**d**) after 15- and 60-min reactions. Gel bands represent the size of the Cy3-labeled primer. Key pausing-positions of interest are defined and color-coded as follows: position **0**, un-extended primer (black); position **8**, primer extension stopped at the pre-displacement site (orange); positions **9-13**, primer stopped at the early-displacement site (blue); positions **16-18**, primer stopped at the mid-PPT displacement site (red); and position **40**, fully extended primer (gray). This color coding is consistently applied to the stacked bar graphs below the gels, which represent the quantification of these bands from three independent experiments. Statistical significance was determined using a one-way ANOVA with Dunnett’s multiple comparisons test, comparing each mutant to the WT control on the corresponding gel. *P < 0.05; **P < 0.01; ***P < 0.001; ****P < 0.0001; bars without asterisks are not significant (P > 0.05).

**Supplementary Fig. 8.**
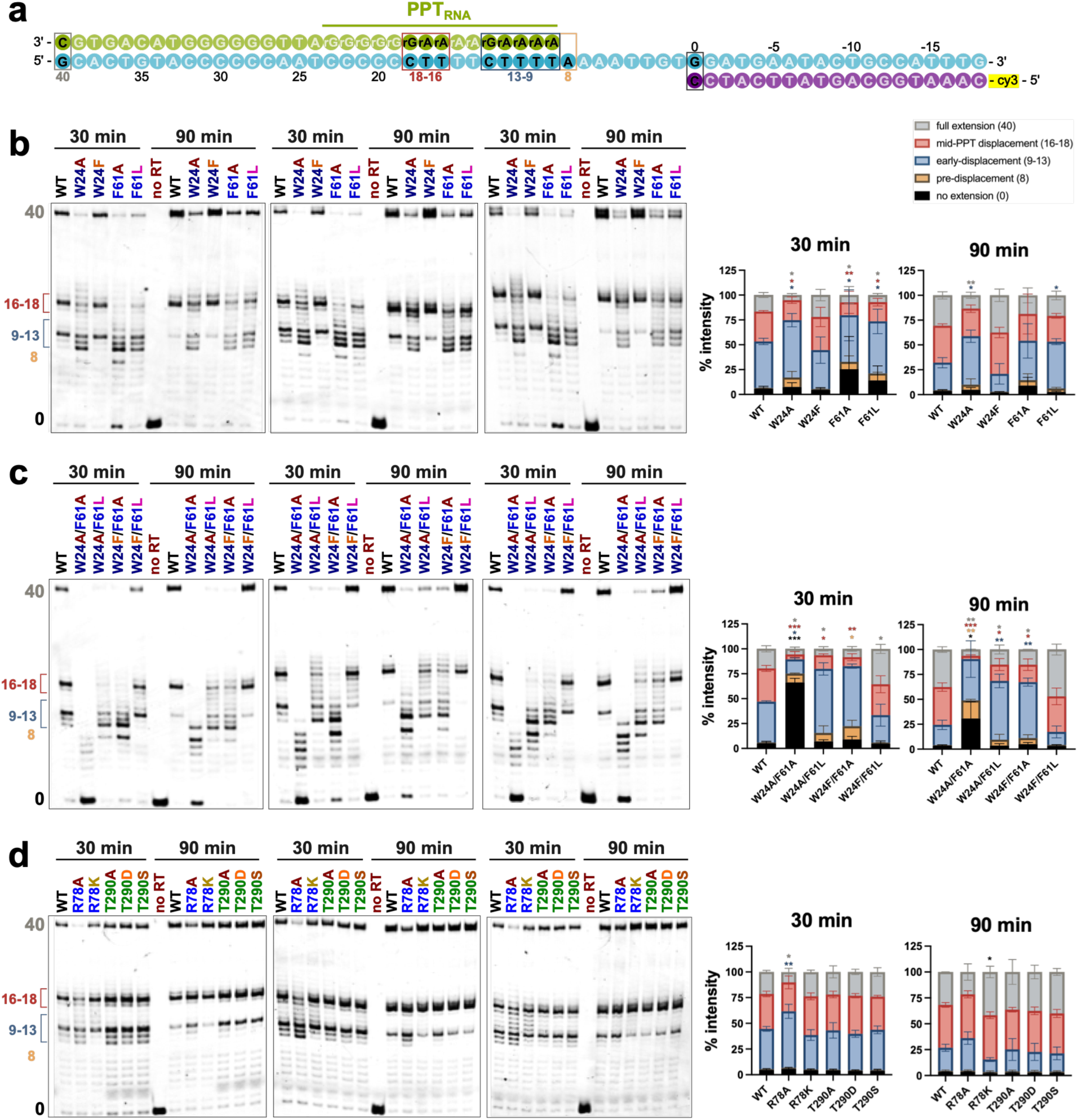
Triplicate primer-extension assays using the PPT_RNA_ displacement (T-P-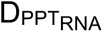) substrate. **a** Schematic of the T-P-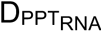 nucleic acid substrate. The primer strand (purple) is labeled with a Cy3 fluorophore at the 5’-end, the template strand is cyan, and PPT_RNA_ displacement strand color in yellow-green. Boxed, darker-colored nucleotides represent strong primer-extension pause sites observed in the assays. **b-d** Triplicate PAGE gels and corresponding quantitative bar graphs for primer-extension assays performed with single W24 and F61 mutants (**b**), double W24 and F61 mutants (**c**), and single R78 and T290 mutants (**d**) after 30- and 90-min reactions. Gel bands represent the size of the Cy3-labeled primer. Key pausing-positions of interest are defined and color-coded as follows: position **0**, un-extended primer (black); position **8**, primer extension stopped at the pre-displacement site (orange); positions **9-13**, primer stopped at the early-displacement site (blue); positions **16-18**, primer stopped at the mid-PPT displacement site (red); and position **40**, fully extended primer (gray). This color coding is consistently applied to the stacked bar graphs below the gels, which represent the quantification of these bands from three independent experiments. Statistical significance was determined using a one-way ANOVA with Dunnett’s multiple comparisons test, comparing each mutant to the WT control on the corresponding gel. *P < 0.05; **P < 0.01; ***P < 0.001; bars without asterisks are not significant (P > 0.05).

**Supplementary Fig. 9.**
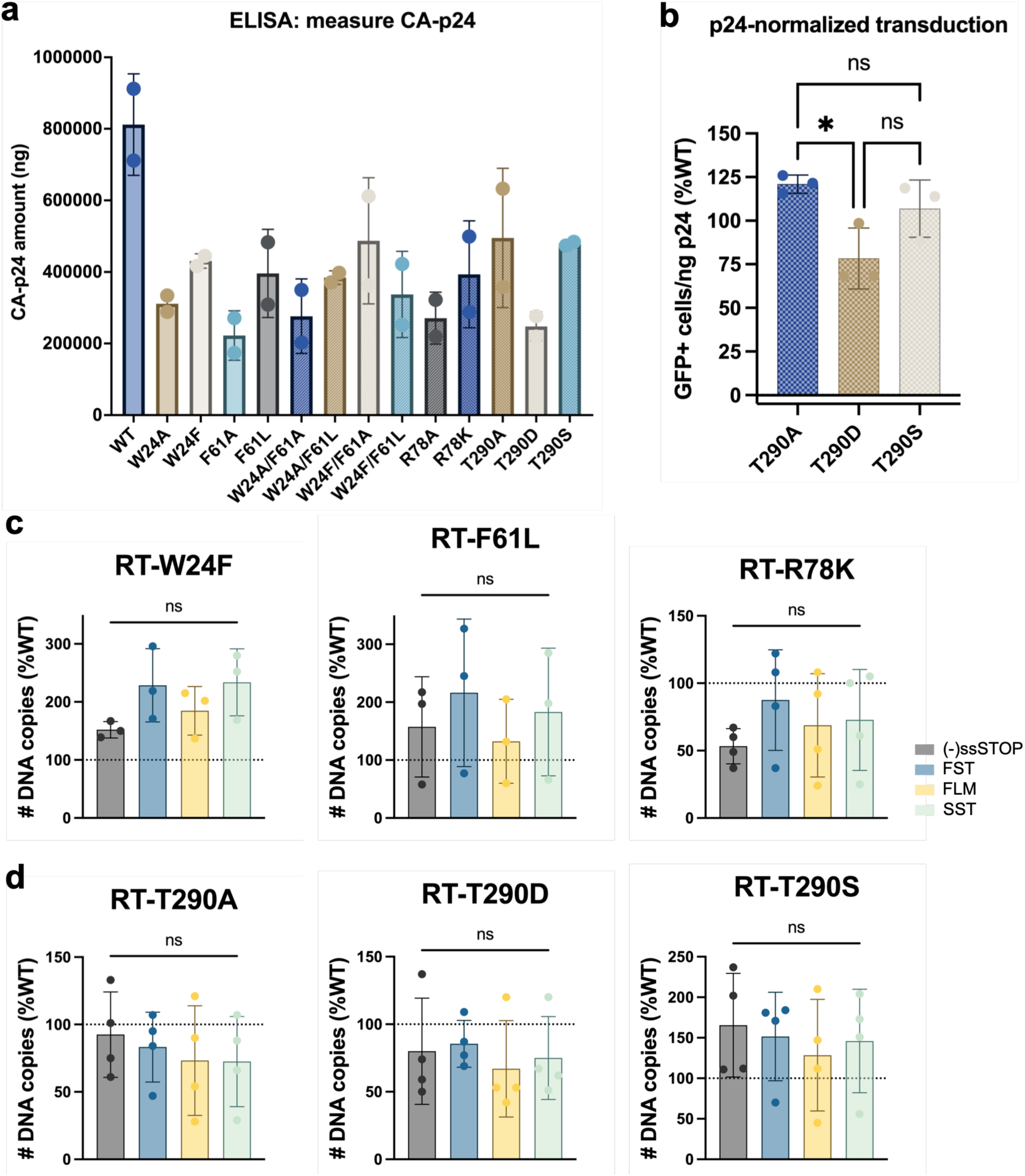
Virological impact of strand displacement rescue and T290 mutants on HIV-1 infection and viral cDNA synthesis. **a** CA-p24 amounts in WT and each RT-mutant pseudotyped virion, as measured by ELISA from two independent experiments. **b** Transduction efficiency of virions carrying RT-T290 mutants in TZM-GFP cells. Statistical significance was determined using a one-way ANOVA with Tukey’s multiple comparisons test, comparing transduction levels to one another. *P < 0.05; ns, not significant. **c-d** Endogenous reverse transcription (ERT) assays of rescue single-mutants (**c**) and RT-T290 mutants (**d**). Bar graphs show the number of DNA copies for products at different stages of reverse transcription from three or four independent experiments ((−)strand strong stop ((−)ssSTOP), first strand transfer (FST), full-length (−)strand (FLM), and second strand transfer (SST)), normalized to the corresponding product level in WT virions. Statistical significance was determined using a one-way ANOVA with Tukey’s multiple comparisons test, comparing the relative levels of each RT product to one another. ns, not significant.

**Supplementary Fig. 10.**
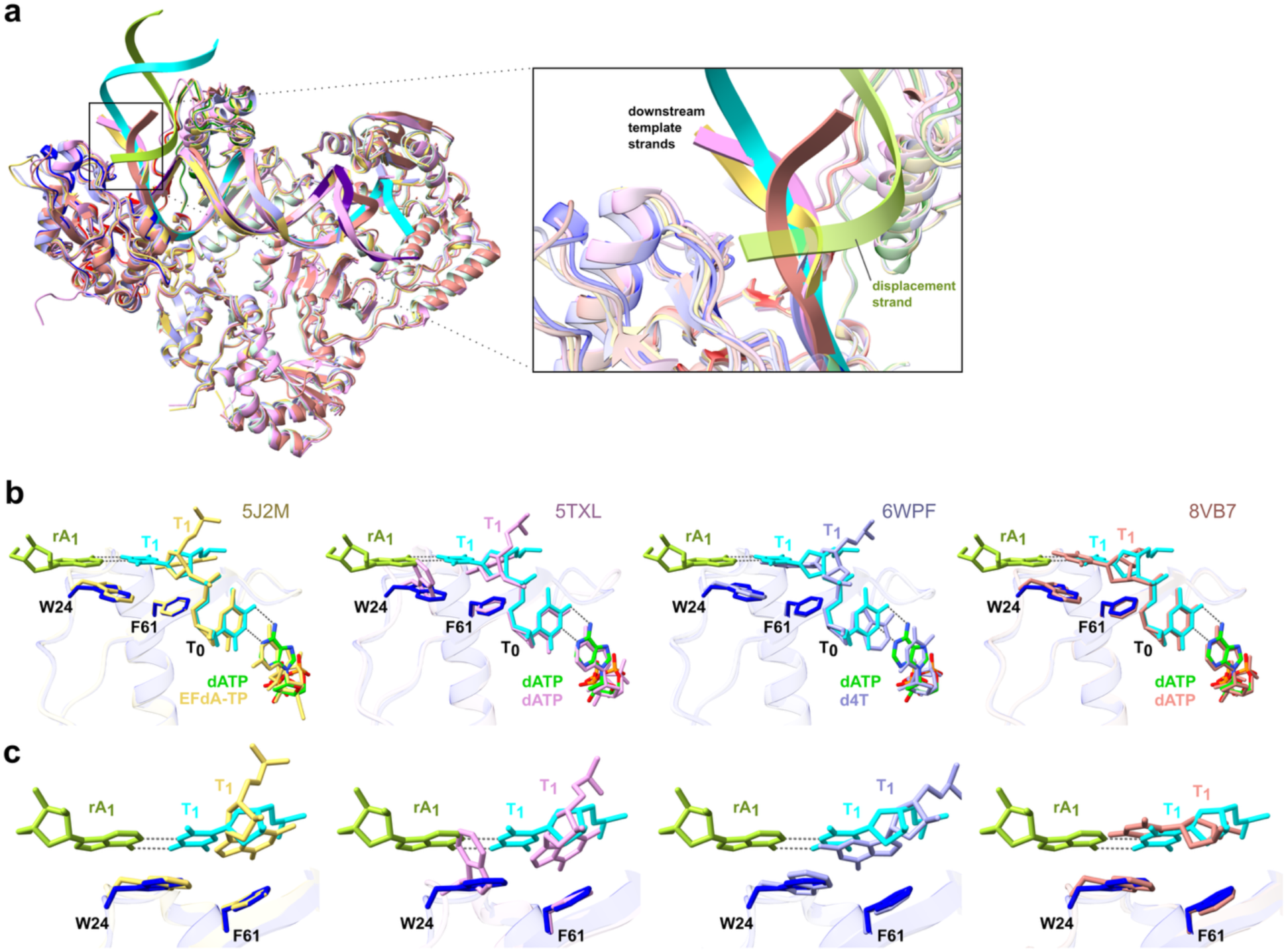
Structural comparison of RT/T-P-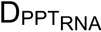/dATP with RT/T-P/nucleotide ternary complexes. **a** Structural superposition of the RT/T-P-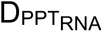/dATP complex with X-ray crystal structures of the RT/T-P/EFdA-TP (PDB: 5J2M; yellow), RT/T-P/dATP (PDB: 5TXL; pink), and RT/T-P/d4T-TP (PDB: 6WPF; light blue) complexes, as well as the cryo-EM structure of the RT/T-P/dATP complex (PDB: 8VB7; pastel red). **Insert:** A zoomed-in view of the template strand overhang site. **b** Structural comparisons of RT/T-P-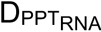/dATP with each of the reference structures listed in a, focusing on the strand-displacement (SD) site and the incoming nucleotide. **c** Zoomed-in views of the corresponding panels in **b** at the SD site, detailing the coordination between specific residues and nucleotides. For all panels, the RT/T-P-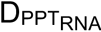/dATP complex is colored as follows: the RT_p66_ fingers subdomain is in blue, the template strand in cyan, the PPT_RNA_ displacement strand in light green, and the incoming dATP in lime.

## Notes

### Competing Interest Statement

The authors have declared no competing interest.

