## Supplementary Figures 1-10 for "Structural Basis of Polypurine Track Strand Displacement by HIV-1 Reverse Transcriptase"

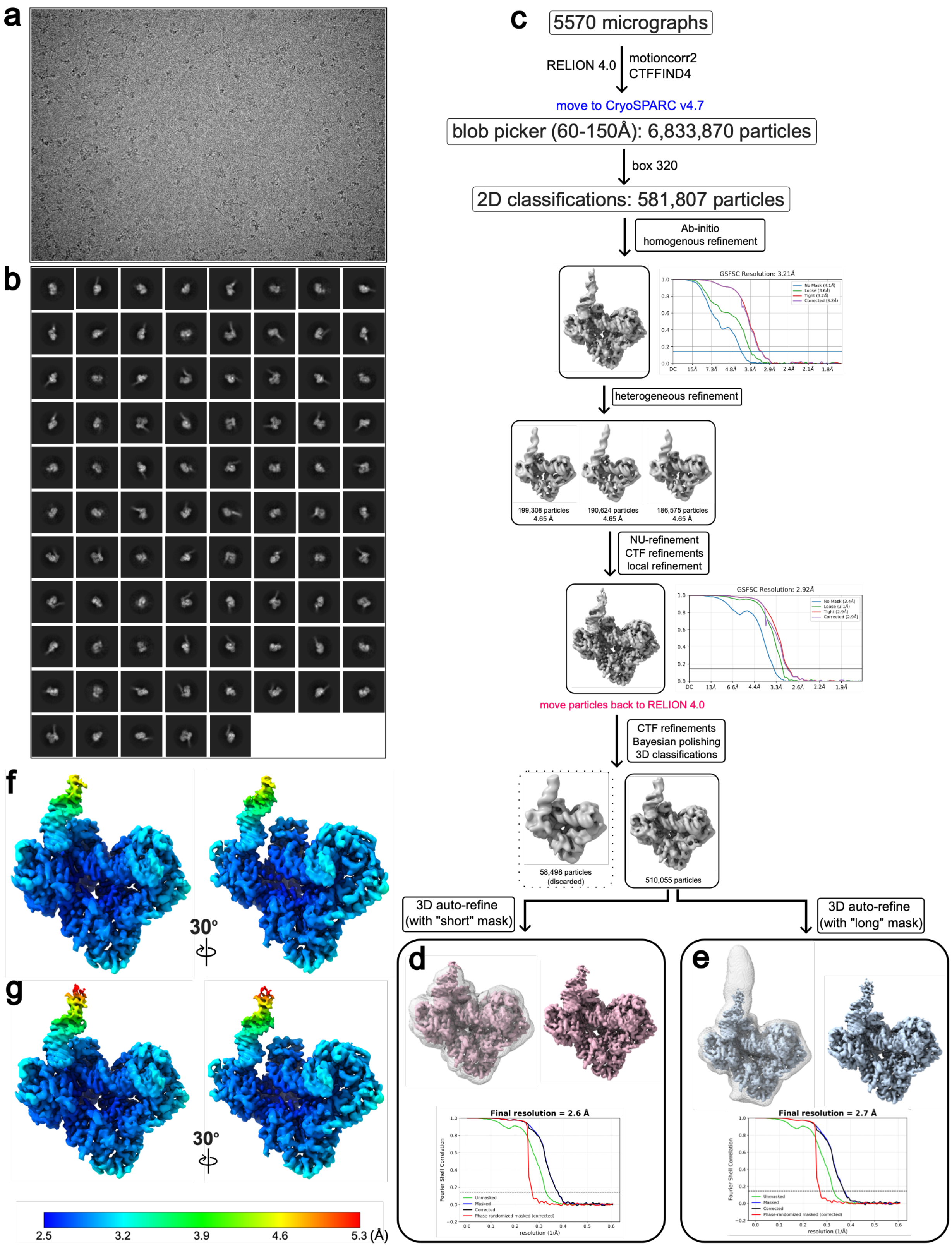

a
b
c
5570 micrographs
RELION 4.0
motioncorr2
CTFFIND4
move to CryoSPARC v4.7
blob picker (60-150Å): 6,833,870 particles
box 320
2D classifications: 581,807 particles
Ab-initio
homogenous refinement
GSFSC Resolution: 3.21Å
heterogeneous refinement
199,308 particles
4.65 Å
190,624 particles
4.65 Å
186,575 particles
4.65 Å
NU-refinement
CTF refinements
local refinement
GSFSC Resolution: 2.92Å
move particles back to RELION 4.0
CTF refinements
Bayesian polishing
3D classifications
58,498 particles
(discarded)
510,055 particles
3D auto-refine
(with "short" mask)
3D auto-refine
(with "long" mask)
d
Final resolution = 2.6 Å
e
Final resolution = 2.7 Å
f
30°
g
30°
2.5
3.2
3.9
4.6
5.3 (Å)

**Supplementary Fig. 1 | Workflow of cryo-EM processing of the RT/T-P-$D_{PPT_{\underline{DNA}}}$/dATP complex.** **a** Representative cryo-EM micrograph. **b** 2D classes. **c** Cryo-EM data processing workflow performed in CryoSPARC v4.7 and RELION 4.0. The number of particles selected at each stage is indicated. **d** Final 3D auto-refined map (unsharpened) contoured at 6.7σ and corresponding Gold-standard Fourier Shell Correlation (FSC) curve using a “short” mask. The final global resolution is 2.6 Å at the FSC = 0.143 criterion. **e** Final 3D auto-refined map (unsharpened) contoured at 6.0σ and corresponding Gold-standard FSC curve using a “long” mask. The final global resolution is 2.7 Å at the FSC = 0.143 criterion. **f-g** Local resolution estimates for the maps shown in **d** and **e**, respectively. The color bar indicates resolution range values in Å.

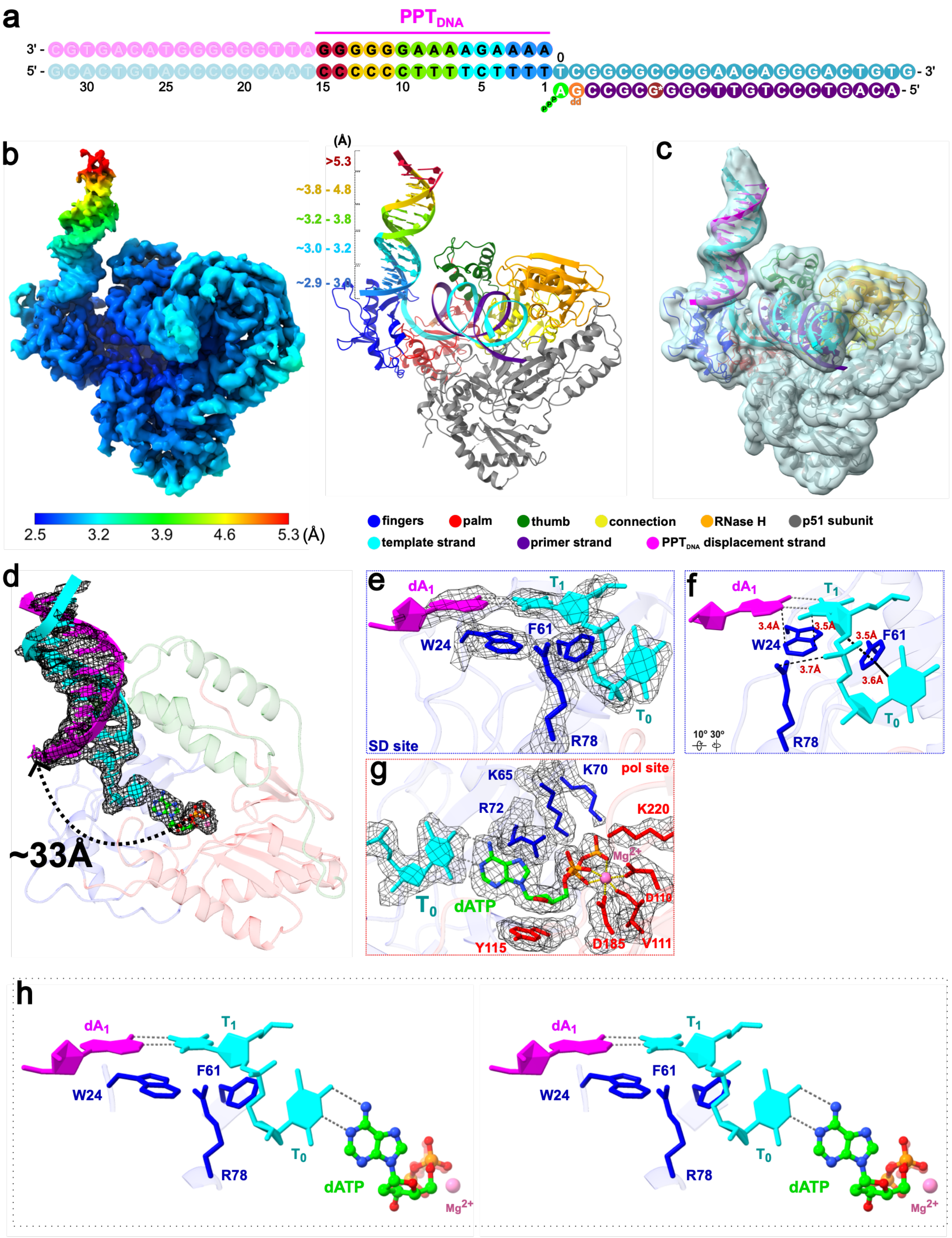
a
PPTDNA
3' -
5' -
0
30
25
20
15
10
5
1
- 3'
- 5'
b
(Å)
>5.3
~3.8 - 4.8
~3.2 - 3.8
~3.0 - 3.2
~2.9 - 3.0
2.5
3.2
3.9
4.6
5.3 (Å)
c
fingers
palm
thumb
connection
RNase H
p51 subunit
template strand
primer strand
PPTDNA displacement strand
d
~33Å
e
dA1
T1
W24
F61
T0
R78
SD site
f
dA1
T1
3.4Å
3.5Å
3.5Å
W24
F61
3.7Å
3.6Å
T0
R78
10° 30°
g
K65
K70
pol site
R72
K220
Mg2+
T0
dATP
D110
Y115
D185
V111
h
dA1
T1
F61
W24
T0
R78
dATP
Mg2+
dA1
T1
F61
W24
T0
R78
dATP
Mg2+

**Supplementary Fig. 2 | Cryo-EM structure of HIV-1 RT in complex with $PPT_{DNA}$ displacement substrate and incoming dATP (RT/T-P-$D_{PPT_{DNA}}$/dATP). a** Schematic of the nucleic acid substrate. The primer strand (purple) contains modifications and coloring as described in **Fig. 2**. The multicolor gradient on the $PPT_{DNA}$ portion of the displacement duplex corresponds to the local resolution range observed in the EM density, as indicated by the color bar in **b**. Faded regions indicate segments not resolved. **b** Cryo-EM density map (**left**) and atomic model (**right**) of the RT/T-P-$D_{PPT_{DNA}}$/dATP complex with the extended 15-bp displacement duplex. The density map was refined using the “long” mask (**Supplementary Fig. 1e**) and is contoured at 6.0σ. Local resolution colors of the displacement duplex are mapped onto the atomic model as indicated in **a**. The color bar indicates resolution range values in Å. **c** Gaussian low-pass filtered map (σ=1.3 Å) contoured at 5.4σ (light blue), was used to guide the modeling of the extended displacement duplex. **d** Close-up view of the downstream displacement duplex and incoming dATP (corresponding to **Fig. 2f**). The cryo-EM density (mesh) from the 3D-refined map (unsharpened) is contoured at 8.1σ. The approximate distance (~33 Å) between the α-phosphate of the incoming dATP and the O5’ atom of the first displacement strand nucleotide (dA1) is indicated. **e** Cryo-EM density (mesh) from the postprocessed map, overlaid on the atomic model of the strand displacement (SD) site shown in **Fig. 2g**. The map is contoured at 10.9σ. **f** View of the SD site rotated relative to **e**. **g** Cryo-EM density (mesh) from the postprocessed map, overlaid on the atomic model of the polymerase active site shown in **Fig. 2h**. The map is contoured at 7.1σ. **h** Stereo view (wall-eyed) of the SD site residues and the incoming dATP. Distances are shown as black dash lines, and hydrogen bonds are shown as gray dash lines.

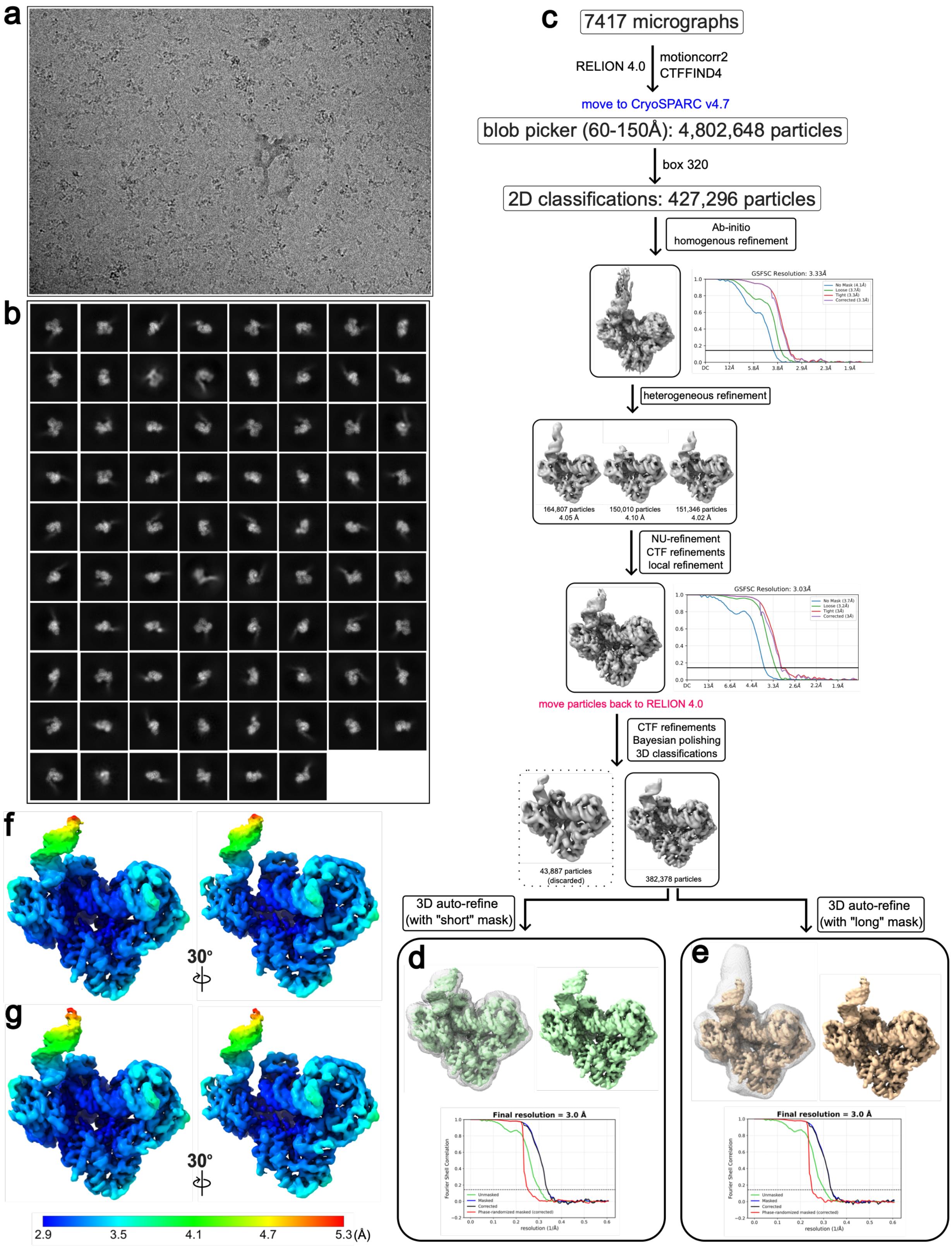
a
b
c
7417 micrographs
RELION 4.0
motioncorr2
CTFFIND4
move to CryoSPARC v4.7
blob picker (60-150Å): 4,802,648 particles
box 320
2D classifications: 427,296 particles
Ab-initio
homogenous refinement
GSFSC Resolution: 3.33Å
heterogeneous refinement
164,807 particles
4.05 Å
150,010 particles
4.10 Å
151,346 particles
4.02 Å
NU-refinement
CTF refinements
local refinement
GSFSC Resolution: 3.03Å
move particles back to RELION 4.0
CTF refinements
Bayesian polishing
3D classifications
43,887 particles
(discarded)
382,378 particles
3D auto-refine
(with "short" mask)
3D auto-refine
(with "long" mask)
d
e
Final resolution = 3.0 Å
Final resolution = 3.0 Å
f
g
30°
30°
2.9
3.5
4.1
4.7
5.3(Å)

**Supplementary Fig. 3 | Workflow of cryo-EM processing of the RT/T-P-$D_{PPT_{\underline{RNA}}}$/dATP complex.** **a** Representative cryo-EM micrograph. **b** 2D classes. **c** Cryo-EM data processing workflow performed in CryoSPARC v4.7 and RELION 4.0. The number of particles selected at each stage is indicated. **d** Final 3D auto-refined map (unsharpened) contoured at 7.4σ and corresponding Gold-standard Fourier Shell Correlation (FSC) curve using a “short” mask. The final global resolution is 3 Å at the FSC = 0.143 criterion. **e** Final 3D auto-refined map (unsharpened) contoured at 6.6σ and corresponding Gold-standard FSC curve using a “long” mask. The final global resolution is 3 Å at the FSC = 0.143 criterion. **f-g** Local resolution estimates for the maps shown in **d** and **e**, respectively. The color bar indicates resolution range values in Å.

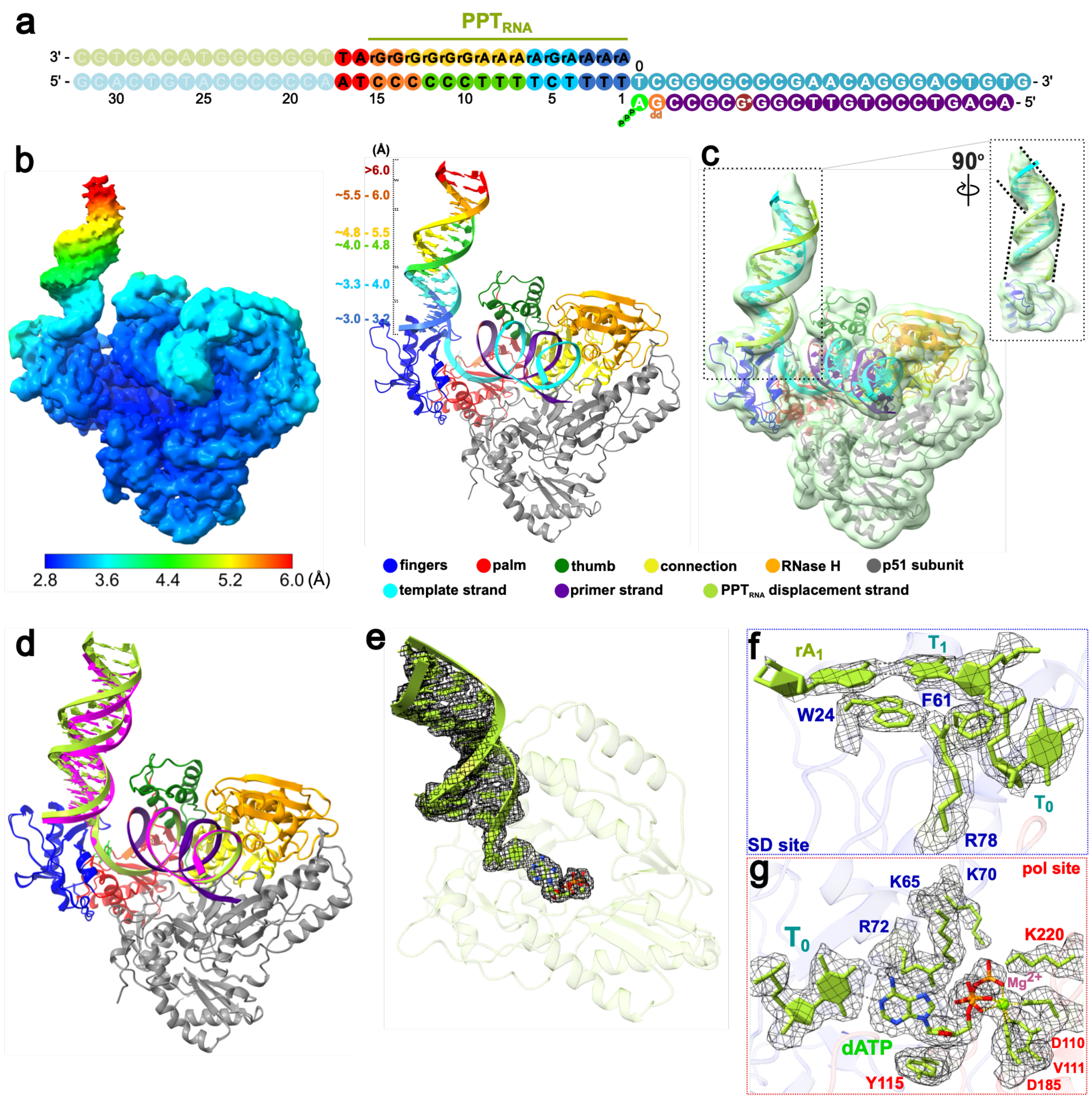

a
PPTRNA
3' -
5' -
- 3'
- 5'
30
25
20
15
10
5
1
0
b
(Å)
>6.0
~5.5 - 6.0
~4.8 - 5.5
~4.0 - 4.8
~3.3 - 4.0
~3.0 - 3.2
2.8
3.6
4.4
5.2
6.0 (Å)
c
90°
fingers
palm
thumb
connection
RNase H
p51 subunit
template strand
primer strand
PPTRNA displacement strand
d
e
f
rA1
T1
W24
F61
T0
SD site
R78
g
K65
K70
pol site
R72
K220
T0
Mg2+
dATP
D110
V111
Y115
D185

**Supplementary Fig. 4 | Cryo-EM structure of HIV-1 RT in complex with $\mathrm{PPT}_{\mathrm{RNA}}$ displacement substrate and incoming dATP (RT/T-P-$\mathrm{D}_{\mathrm{PPT}_{\underline{\mathrm{RNA}}}}$/dATP). a** Schematic of the nucleic acid substrate. The primer strand (purple) contains modifications and coloring as described in **Fig. 2**. The multicolor gradient on the $\mathrm{PPT}_{\mathrm{DNA}}$ portion of the displacement duplex corresponds to the local resolution range observed in the EM density, as indicated by the color bar in **b**. Faded regions indicate segments not resolved. **b** Cryo-EM density map (**left**) and atomic model (**right**) of the RT/T-P-$\mathrm{D}_{\mathrm{PPT}_{\underline{\mathrm{RNA}}}}$/dATP complex with the extended 17-bp displacement duplex. The density map was refined using the “long” mask (**Supplementary Fig. 2e**) and is contoured at 5.2σ. Local resolution colors of the displacement duplex are mapped onto the atomic model as indicated in **a**. The color bar indicates resolution range values in Å. **c** Gaussian low-pass filtered map (σ=1.3 Å; light green), contoured at 5.4σ, used to guide modeling of the extended 17-bp $\mathrm{PPT}_{\mathrm{RNA}}$ displacement duplex. Right: a 90° rotated view of the $\mathrm{PPT}_{\mathrm{RNA}}$ displacement duplex, with black dashed lines indicating its helical axis directions. An apparent bend occurs within the region of consecutive rG of the $\mathrm{PPT}_{\mathrm{RNA}}$ sequence. **d** Structural overlay of the extended (15-bp) RT/T-P-$\mathrm{D}_{\mathrm{PPT}_{\underline{\mathrm{DNA}}}}$/dATP and extended (17-bp) RT/T-P-$\mathrm{D}_{\mathrm{PPT}_{\underline{\mathrm{RNA}}}}$/dATP complexes. Enzyme and primer-strand are colored as described previously. The template and displacement strands are colored magenta for the $\mathrm{PPT}_{\mathrm{DNA}}$ model and yellow-green for the $\mathrm{PPT}_{\mathrm{RNA}}$ model. **e** Close-up view of the downstream displacement duplex and incoming dATP (corresponding to the view in **Fig. 3f**). The cryo-EM density (mesh) from the 3D-refined map (unsharpened) is contoured at 7.4σ. **f** Cryo-EM density (mesh) from the post-processed map, overlaid on the atomic model of the strand displacement (SD) site shown in **Fig. 3g**. The map is contoured at 7.2σ. **g** Cryo-EM density (mesh) from the post-processed map, overlaid on the atomic model of the polymerase (pol) active site shown in **Fig. 3h**. The map is contoured at 5σ. Dashed gray lines indicate hydrogen bonds.

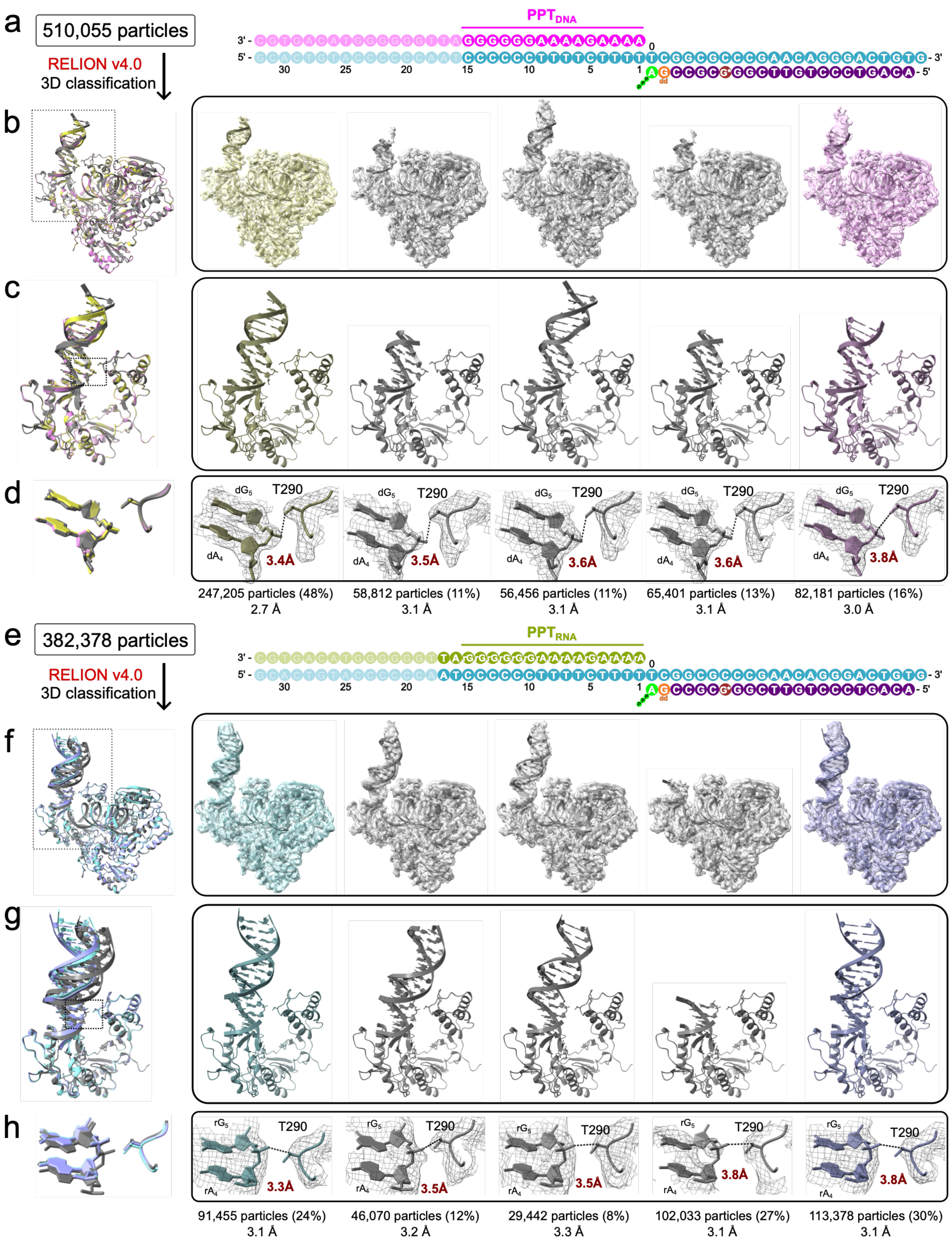

a
510,055 particles
RELION v4.0
3D classification
PPT_DNA
b
c
d
dG5
T290
dA4
3.4Å
3.5Å
3.6Å
3.6Å
3.8Å
247,205 particles (48%)
2.7 Å
58,812 particles (11%)
3.1 Å
56,456 particles (11%)
3.1 Å
65,401 particles (13%)
3.1 Å
82,181 particles (16%)
3.0 Å
e
382,378 particles
RELION v4.0
3D classification
PPT_RNA
f
g
h
rG5
T290
rA4
3.3Å
3.5Å
3.5Å
3.8Å
3.8Å
91,455 particles (24%)
3.1 Å
46,070 particles (12%)
3.2 Å
29,442 particles (8%)
3.3 Å
102,033 particles (27%)
3.1 Å
113,378 particles (30%)
3.1 Å

**Supplementary Fig. 5 | 3D classifications of the final particle stacks for the RT/T-P-$\mathrm{D_{PPT_{DNA}}}$/dATP and RT/T-P-$\mathrm{D_{PPT_{RNA}}}$/dATP complexes. a-d** 3D classification (5 classes) of the final particle stack for the RT/T-P-$\mathrm{D_{PPT_{DNA}}}$/dATP complex. **e-h** 3D classification (5 classes) of the final particle stack for the RT/T-P-$\mathrm{D_{PPT_{RNA}}}$/dATP complex. **a, e** Left: number of particles used for 3D classification. Right: schematic of the nucleic acid substrate. **b, f** Overlay of the atomic models for the five 3D classes. **Right panels**: Individual classes showing atomic models fitted into their corresponding unsharpened 3D-refined maps. All maps are shown at an absolute contour level of 0.0055. **c, g** Close-up view (boxed region in **b, f**) showing the overlay of the polymerase subdomain and the downstream displacement duplex. **Right panels**: Individual classes at this view. **d, h** Close-up view (boxed region in **c, g**) detailing the contact between T290 and the displacement strand. **Right panels**: Individual classes at this view, with the distance between T290 and the $\mathrm{dG_5}$ (**d**) or $\mathrm{rG_5}$ (**h**) phosphate indicated for each. The corresponding unsharpened 3D-refined density maps (mesh) are shown at an absolute contour level of 0.0055 (except 0.0035 for class 4 in **h**). The class exhibiting the closest T290 contact with the $\mathrm{PPT_{DNA}}$ (PDB 11XI, EMDB 76156) is colored yellow (**d**); the closest T290 contact with the $\mathrm{PPT_{RNA}}$ (PDB 11XM, EMDB 76160) is colored light blue (**h**). The class with the furthest T290-$\mathrm{PPT_{DNA}}$ contact (PDB 11XJ, EMDB 76157) is colored pink (**d**); the furthest T290-$\mathrm{PPT_{DNA}}$ contact (PDB 11XN, EMDB 76161) is colored light purple (**h**). All other classes with intermediate contacts are gray. The number of particles, percentage of the total stack, and final map resolution are indicated for each class.

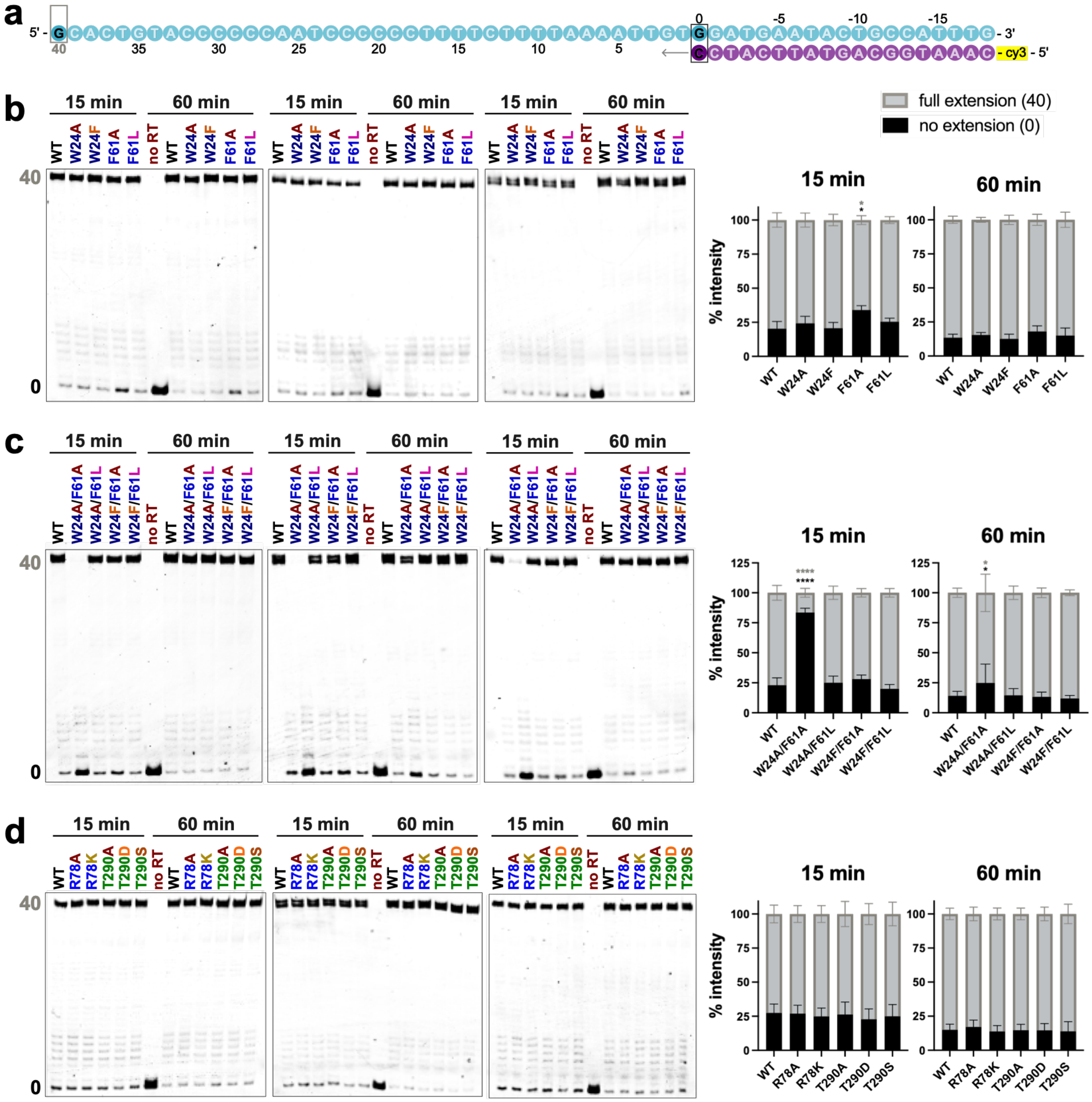


**Supplementary Fig. 6 | Triplicate primer-extension assays using the template-primer (T-P) only substrate. a** Schematic of the T-P only nucleic acid substrate. The primer strand (purple) is labeled with a Cy3 fluorophore at the 5'-end, and the template strand is cyan. Boxed, darker-colored nucleotides represent strong primer-extension pause sites observed in the assays. **b-d** Triplicate PAGE gels and corresponding quantitative bar graphs for primer-extension assays performed with single W24 and F61 mutants (**b**), double W24 and F61 mutants (**c**), and single R78 and T290 mutants (**d**) after 15- and 60-min reactions. Gel bands represent the size of the Cy3-labeled primer. Quantified positions of interest include the un-extended primer at position 0 (black) and the fully extended primer at position 40 (gray). Statistical significance was determined using a one-way ANOVA with Dunnett's multiple comparisons test, comparing each mutant to the WT control on the corresponding gels. P < 0.05; ****P < 0.0001; bars without asterisks are not significant (P > 0.05).

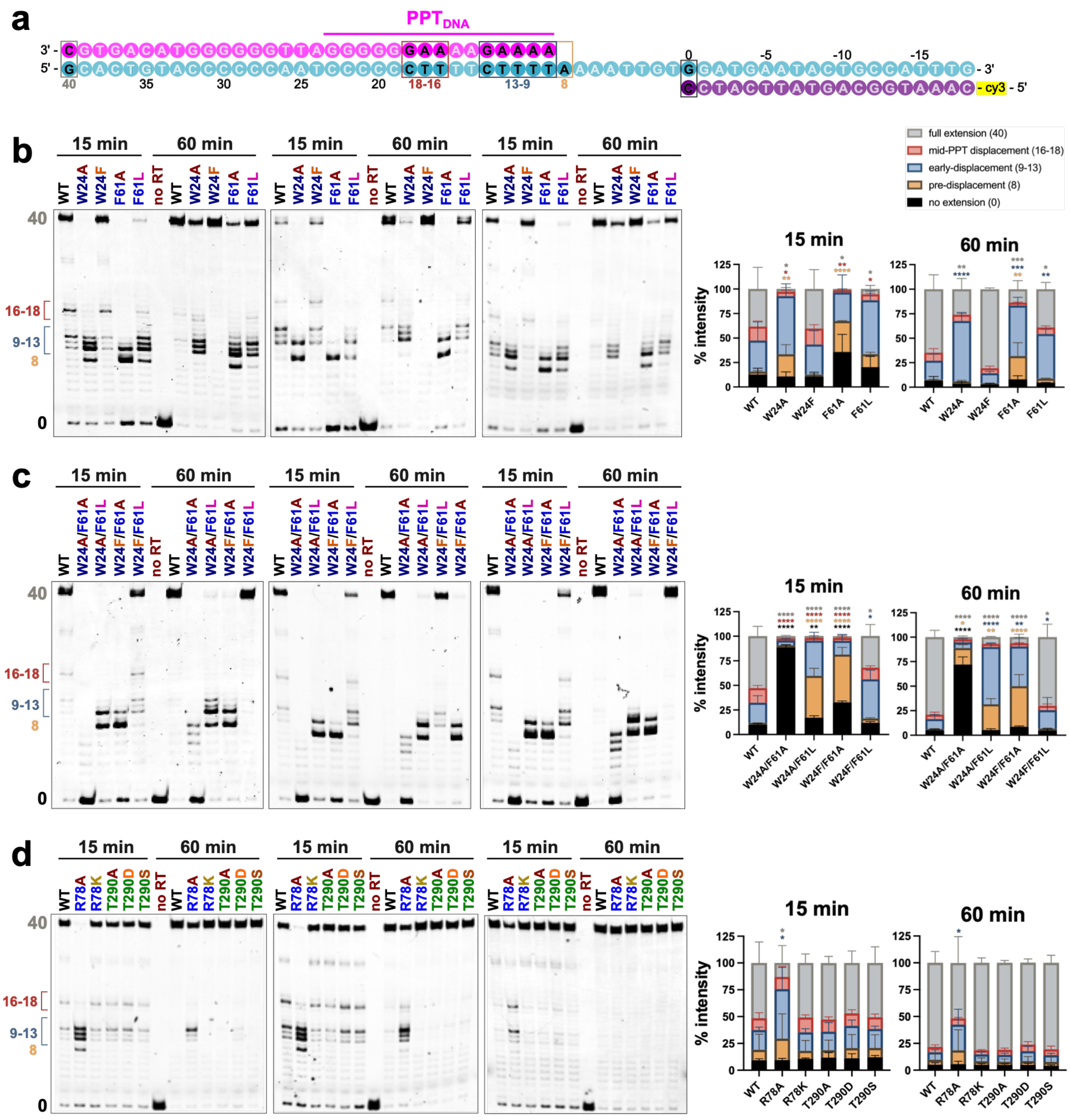


**Supplementary Fig. 7 | Triplicate primer-extension assays using the $PPT_{DNA}$ displacement (T-P-$D_{PPT_{DNA}}$) substrate. a** Schematic of the T-P-$D_{PPT_{DNA}}$ nucleic acid substrate. The primer strand (purple) is labeled with a Cy3 fluorophore at the 5'-end, the template strand is cyan, and $PPT_{DNA}$ displacement strand color in magenta. Boxed, darker-colored nucleotides represent strong primer-extension pause sites observed in the assays. **b-d** Triplicate PAGE gels and corresponding quantitative bar graphs for primer-extension assays performed with single W24 and F61 mutants (**b**), double W24 and F61 mutants (**c**), and single R78 and T290 mutants (**d**) after 15- and 60-min reactions. Gel bands represent the size of the Cy3-labeled primer. Key pausing-positions of interest are defined and color-coded as follows: position **0**, un-extended primer (black); position **8**, primer extension stopped at the pre-displacement site (orange); positions **9-13**, primer stopped at the early-displacement site (blue); positions **16-18**, primer stopped at the mid-PPT displacement site (red); and position **40**, fully extended primer (gray). This color coding is consistently applied to the stacked bar graphs below the gels, which represent the quantification of these bands from three independent experiments. Statistical significance was determined using a one-way ANOVA with Dunnett's multiple comparisons test, comparing each mutant to the WT control on the corresponding gel. *$P < 0.05$; **$P < 0.01$; ***$P < 0.001$; ****$P < 0.0001$; bars without asterisks are not significant ($P > 0.05$).

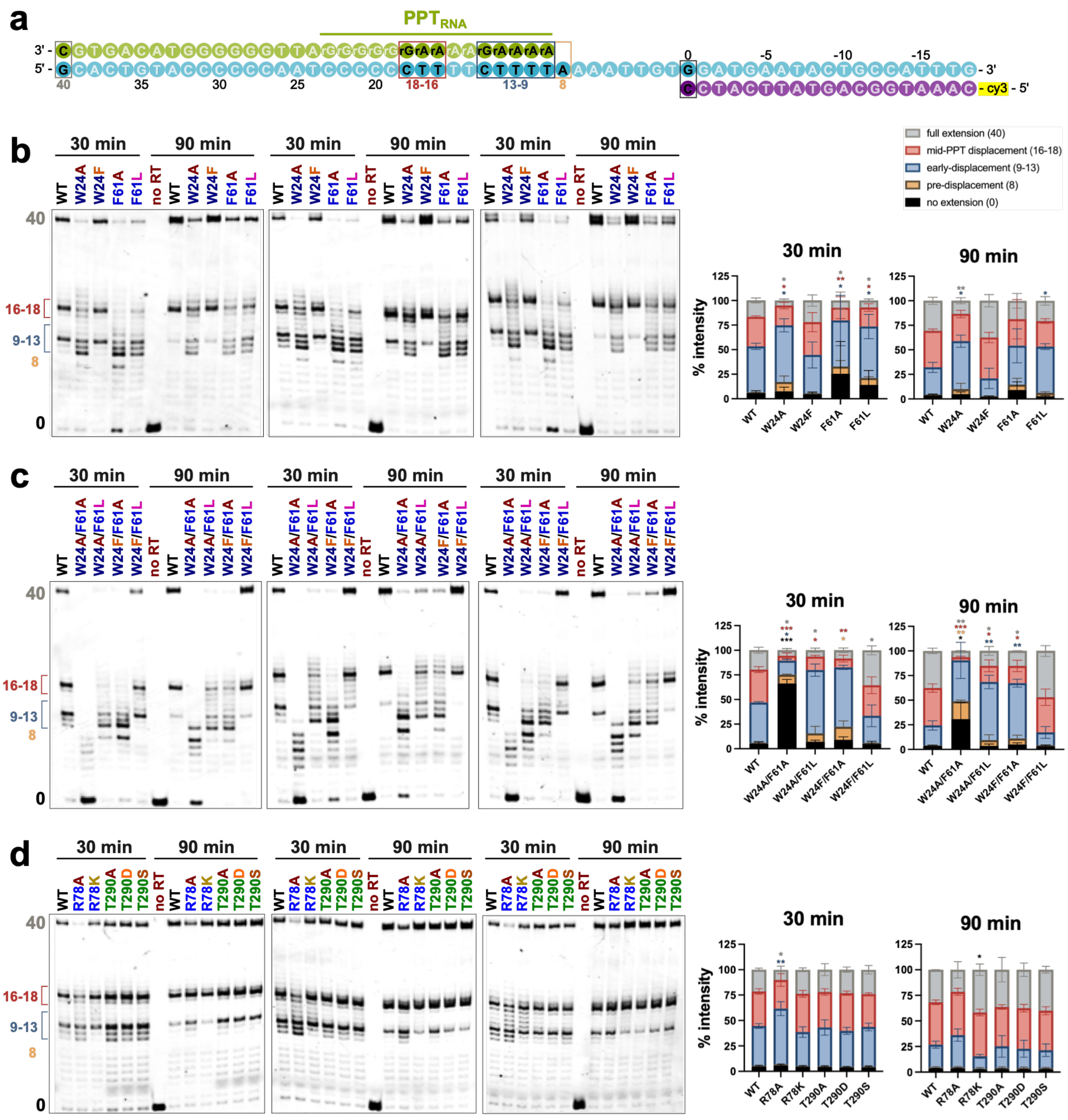


**Supplementary Fig. 8 | Triplicate primer-extension assays using the PPT$_{RNA}$ displacement (T-P-D$_{PPT_{RNA}}$) substrate. a** Schematic of the T-P-D$_{PPT_{RNA}}$ nucleic acid substrate. The primer strand (purple) is labeled with a Cy3 fluorophore at the 5'-end, the template strand is cyan, and PPT$_{RNA}$ displacement strand color in yellow-green. Boxed, darker-colored nucleotides represent strong primer-extension pause sites observed in the assays. **b-d** Triplicate PAGE gels and corresponding quantitative bar graphs for primer-extension assays performed with single W24 and F61 mutants (**b**), double W24 and F61 mutants (**c**), and single R78 and T290 mutants (**d**) after 30- and 90-min reactions. Gel bands represent the size of the Cy3-labeled primer. Key pausing-positions of interest are defined and color-coded as follows: position **0**, un-extended primer (black); position **8**, primer extension stopped at the pre-displacement site (orange); positions **9-13**, primer stopped at the early-displacement site (blue); positions **16-18**, primer stopped at the mid-PPT displacement site (red); and position **40**, fully extended primer (gray). This color coding is consistently applied to the stacked bar graphs below the gels, which represent the quantification of these bands from three independent experiments. Statistical significance was determined using a one-way ANOVA with Dunnett's multiple comparisons test, comparing each mutant to the WT control on the corresponding gel. *P < 0.05; **P < 0.01; ***P < 0.001; bars without asterisks are not significant (P > 0.05).

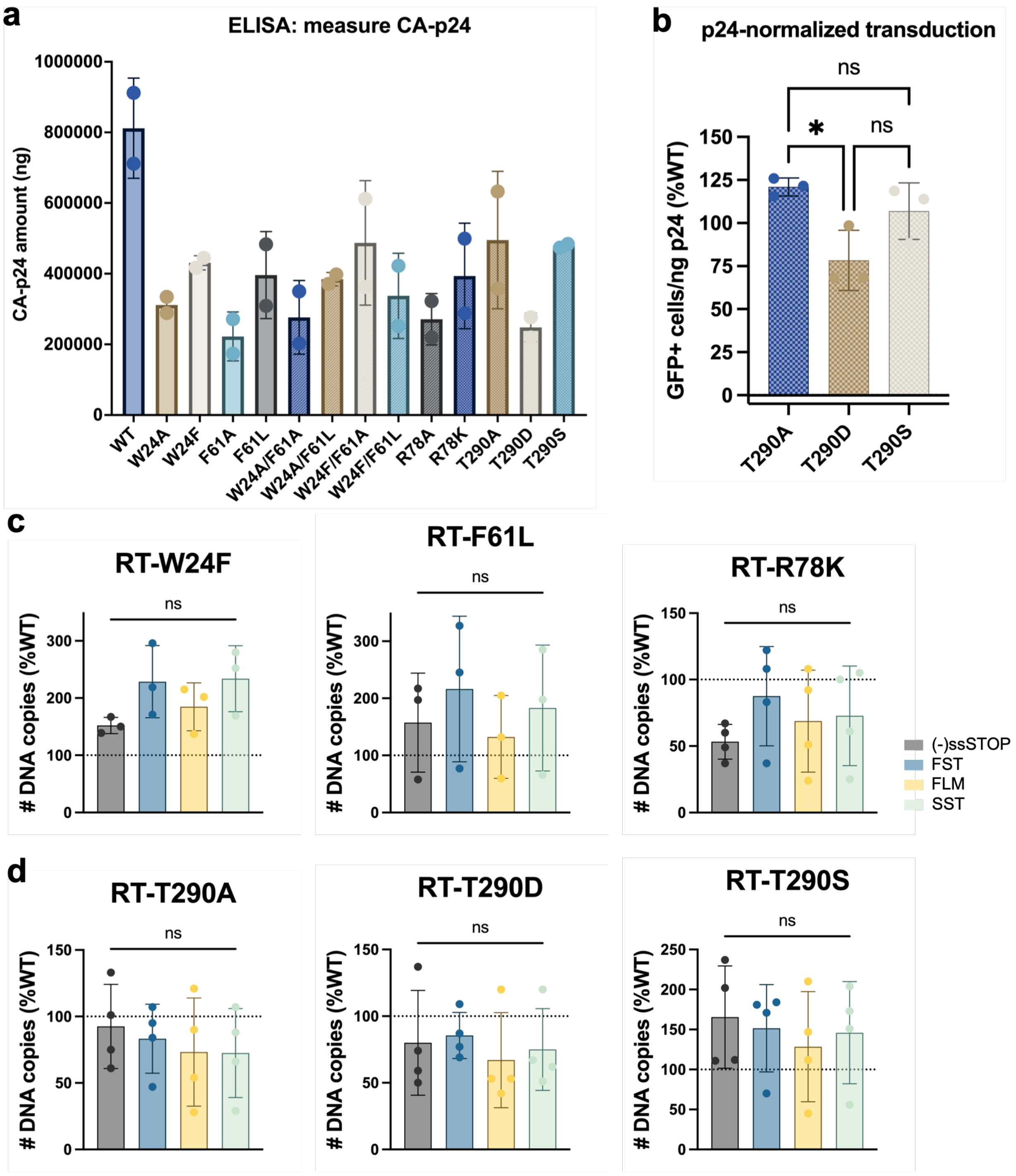
a
ELISA: measure CA-p24
CA-p24 amount (ng)
1000000
800000
600000
400000
200000
0
WT
W24A
W24F
F61A
F61L
W24A/F61A
W24A/F61L
W24F/F61A
W24F/F61L
R78A
R78K
T290A
T290D
T290S
b
p24-normalized transduction
GFP+ cells/ng p24 (%WT)
150
125
100
75
50
25
0
ns
*
ns
T290A
T290D
T290S
c
RT-W24F
RT-F61L
RT-R78K
### DNA copies (%WT)
ns
(-)ssSTOP
FST
FLM
SST
d
RT-T290A
RT-T290D
RT-T290S
### DNA copies (%WT)
ns

**Supplementary Fig. 9 | Virological impact of strand displacement rescue and T290 mutants on HIV-1 infection and viral cDNA synthesis. a** CA-p24 amounts in WT and each RT-mutant pseudotyped virion, as measured by ELISA from two independent experiments. **b** Transduction efficiency of virions carrying RT-T290 mutants in TZM-GFP cells. Statistical significance was determined using a one-way ANOVA with Tukey's multiple comparisons test, comparing transduction levels to one another. *$P < 0.05$; ns, not significant. **c-d** Endogenous reverse transcription (ERT) assays of rescue single-mutants (**c**) and RT-T290 mutants (**d**). Bar graphs show the number of DNA copies for products at different stages of reverse transcription from three or four independent experiments ((-)strand strong stop ((-)ssSTOP), first strand transfer (FST), full-length (-)strand (FLM), and second strand transfer (SST)), normalized to the corresponding product level in WT virions. Statistical significance was determined using a one-way ANOVA with Tukey's multiple comparisons test, comparing the relative levels of each RT product to one another. ns, not significant.

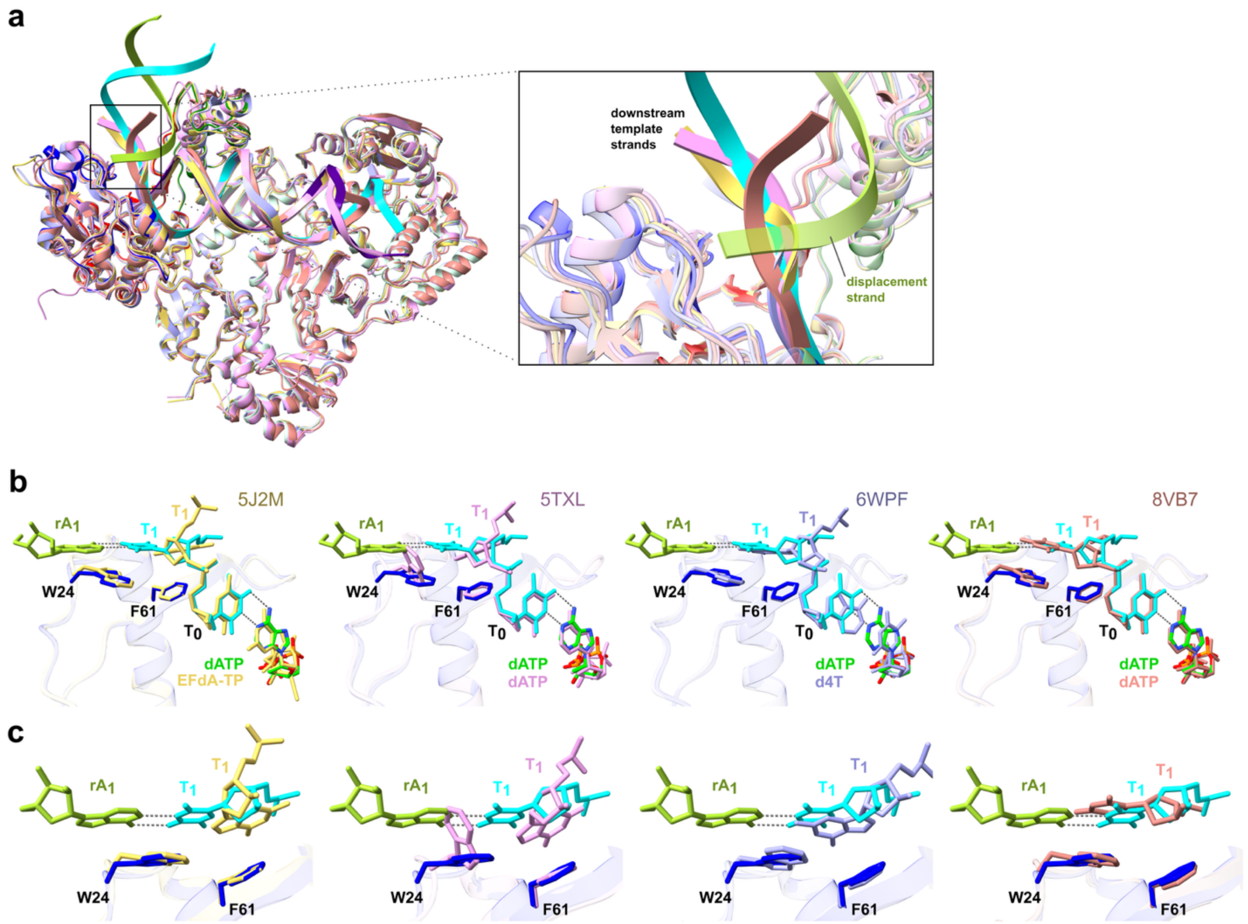


**Supplementary Fig. 10 | Structural comparison of RT/T-P-$D_{PPT_{\underline{RNA}}}$/dATP with RT/T-P/nucleotide ternary complexes. a** Structural superposition of the RT/T-P-$D_{PPT_{\underline{RNA}}}$/dATP complex with X-ray crystal structures of the RT/T-P/EFdA-TP (PDB: 5J2M; yellow), RT/T-P/dATP (PDB: 5TXL; pink), and RT/T-P/d4T-TP (PDB: 6WPF; light blue) complexes, as well as the cryo-EM structure of the RT/T-P/dATP complex (PDB: 8VB7; pastel red). **Insert:** A zoomed-in view of the template strand overhang site. **b** Structural comparisons of RT/T-P-$D_{PPT_{\underline{RNA}}}$/dATP with each of the reference structures listed in a, focusing on the strand-displacement (SD) site and the incoming nucleotide. **c** Zoomed-in views of the corresponding panels in **b** at the SD site, detailing the coordination between specific residues and nucleotides. For all panels, the RT/T-P-$D_{PPT_{\underline{RNA}}}$/dATP complex is colored as follows: the $RT_{p66}$ fingers subdomain is in blue, the template strand in cyan, the $PPT_{RNA}$ displacement strand in light green, and the incoming dATP in lime.
