## Supplementary Tables 1-3 for "Structural Basis of Polypurine Track Strand Displacement by HIV-1 Reverse Transcriptase"

**Supplementary Table 1: Cryo-EM data collection, refinement and validation statistics**

| | RT/T-P-D$_{DNA-PPT}$/dATP (10D) (EMDB-76154) (PDB 11XG) | RT/T-P-D$_{DNA-PPT}$/dATP (15D) (EMDB-76155) (PDB 11XH) | RT/T-P-D$_{DNA-PPT}$/dATP (T290-PPT$_{DNA}$ contact) (EMDB-76156) (PDB 11XI) | RT/T-P-D$_{DNA-PPT}$/dATP (no T290-PPT$_{DNA}$ contact) (EMDB-76157) (PDB 11XJ) | RT/T-P-D$_{RNA-PPT}$/dATP (10D) (EMDB-76158) (PDB 11XK) | RT/T-P-D$_{RNA-PPT}$/dATP (17D) (EMDB-76159) (PDB 11XL) | RT/T-P-D$_{RNA-PPT}$/dATP (T290-PPT$_{RNA}$ contact) (EMDB-76160) (PDB 11XM) | RT/T-P-D$_{RNA-PPT}$/dATP (no T290-PPT$_{RNA}$ contact) (EMDB-76161) (PDB 11XN) |
|---|---|---|---|---|---|---|---|---|
| **Collection and processing** | | | | | | | | |
| Magnification | 105,000 | 105,000 | 105,000 | 105,000 | 105,000 | 105,000 | 105,000 | 105,000 |
| Voltage (kV) | 300 | 300 | 300 | 300 | 300 | 300 | 300 | 300 |
| Electron exposure (e–/Å$^2$) | 50 | 50 | 50 | 50 | 50 | 50 | 50 | 50 |
| Defocus range (μm) | -0.5 to -2.0 | -0.5 to -2.0 | -0.5 to -2.0 | -0.5 to -2.0 | -0.5 to -2.0 | -0.5 to -2.0 | -0.5 to -2.0 | -0.5 to -2.0 |
| Pixel size (Å) | 0.825 | 0.825 | 0.825 | 0.825 | 0.825 | 0.825 | 0.825 | 0.825 |
| Symmetry imposed | C1 | C1 | C1 | C1 | C1 | C1 | C1 | C1 |
| Initial particle images (no.) | 6,833,870 | 6,833,870 | 6,833,870 | 6,833,870 | 4,802,648 | 4,802,648 | 4,802,648 | 4,802,648 |
| Final particle images (no.) | 510,055 | 510,055 | 247,205 | 82,181 | 382,378 | 382,378 | 91,455 | 113,378 |
| Map resolution (Å)<br>FSC threshold | 2.6 | 2.7 | 2.7 | 3.0 | 3.0 | 3.0 | 3.1 | 3.1 |
| **Refinement** | | | | | | | | |
| Initial model used (PDBID) | 5J2M | 11XG | 11XH | 11XG | 11XG | 11XK | 11XL | 11XL |
| Model resolution (Å)<br>FSC threshold | 2.4<br>0.143 | 2.4<br>0.143 | 2.4<br>0.143 | 2.6<br>0.143 | 2.7<br>0.143 | 2.7<br>0.143 | 2.9<br>0.143 | 2.8<br>0.143 |
| Model composition<br>Non-hydrogen atoms<br>Residues: Protein – Nucleotide<br>Ligands | 9263<br>967 – 64<br>MG: 2<br>DTP: 1 | 9468<br>967 – 74<br>MG: 2<br>DTP: 1 | 9386<br>967 – 70<br>MG: 2<br>DTP: 1 | 9263<br>967 – 64<br>MG: 2<br>DTP: 1 | 9273<br>967 – 64<br>MG: 2<br>DTP: 1 | 9565<br>967 – 78<br>MG: 2<br>DTP: 1 | 9552<br>967 – 78<br>MG: 2<br>DTP: 1 | 9562<br>967 – 78<br>MG: 2<br>DTP: 1 |
| *B* factors min/max/mean (Å$^2$)<br>Protein<br>Nucleotide<br>Ligand | 56.4/184.9/98.1<br>63.2/268.9/141.1<br>69.6/110.2/82.4 | 48.2/180.4/88.3<br>52.1/368.1/164.5<br>61.6/96.0/71.7 | 26.3/161.1/63.9<br>31.3/283.3/125.2<br>39.9/69.3/49.4 | 56.6/179.4/92.0<br>67.7/250.6/138.4<br>67.9/103.6/77.0 | 74.9/209.3/118.0<br>90.8/324.5/179.0<br>93.0/132.2/105.7 | 74.7/201.6/119.2<br>91.9/553.2/244.2<br>91.5/133.8/106.7 | 74.7/201.6/119.2<br>91.9/553.2/244.2<br>91.5/133.8/106.7 | 73.2/230.8/122.5<br>94.7/326.9/194.1<br>96.7/138.4/111.1 |
| R.m.s. deviations<br>Bond lengths (Å)<br>Bond angles (°) | 0.006<br>0.513 | 0.005<br>0.511 | 0.006<br>0.532 | 0.005<br>0.531 | 0.005<br>0.534 | 0.004<br>0.545 | 0.003<br>0.507 | 0.003<br>0.493 |
| Validation<br>MolProbity score<br>Clashscore<br>Rotamers outliers (%) | 1.20<br>4.12<br>0.23 | 1.27<br>5.14<br>0.00 | 1.16<br>3.69<br>0.46 | 1.32<br>5.90<br>0.23 | 1.42<br>7.73<br>0.12 | 1.55<br>10.85<br>0.23 | 1.56<br>9.13<br>1.04 | 1.51<br>9.12<br>0.23 |
| Ramachandran plot<br>Favored (%)<br>Allowed (%)<br>Ourliers (%) | 98.12<br>1.88<br>0.00 | 98.12<br>1.88<br>0.00 | 98.23<br>1.77<br>0.00 | 98.33<br>1.67<br>0.00 | 98.33<br>1.67<br>0.00 | 98.33<br>1.67<br>0.00 | 97.70<br>2.30<br>0.00 | 97.91<br>2.09<br>0.00 |

| Mutant | Primers (5' to 3') |
|---|---|
| W24A | F: TGGCCCAAAAGTTAAACAAGCGCCATTGACAGAA<br>R: TTGTTTAACTTTTGGGCCATCCATCCCGG |
| W24F | F: TGGCCCAAAAGTTAAACAATTTCCATTGACAGAA<br>R: TCCATCCCGGGCTTTAATTTTACTGGTACAGTCT |
| F61A | F: TCCATACAATACTCCAGTAGCTGCCATAAAGAAAAAAGACAG<br>R: TACTGGAGTATTGTATGGATTTTCAGGC |
| F61L | F: TCCATACAATACTCCAGTATTGGCCATAAAGAAAAAAGACAG<br>R: TTTTCAGGCCCAATTTTTGAAATTTTCCCTTCC |
| W24A/F61A (from W24A) | F: TCCATACAATACTCCAGTAGCTGCCATAAAGAAAAAAGACAG<br>R: TACTGGAGTATTGTATGGATTTTCAGGC |
| W24A/F61L (from W24A) | F: TCCATACAATACTCCAGTATTGGCCATAAAGAAAAAAGACAG<br>R: TTTTCAGGCCCAATTTTTGAAATTTTCCCTTCC |
| W24F/F61A (from W24F) | F: TCCATACAATACTCCAGTAGCTGCCATAAAGAAAAAAGACAG<br>R: TACTGGAGTATTGTATGGATTTTCAGGC |
| W24F/F61L (from W24F) | F: TCCATACAATACTCCAGTATTGGCCATAAAGAAAAAAGACAG<br>R: TTTTCAGGCCCAATTTTTGAAATTTTCCCTTCC |
| R78A | F: AGTAGATTTCGCAGAACTTAATAAGAGAACTCAAGACTTCTG<br>R: AATTTTCTCCATTTAGTACTGTCTTTTTTCTTTATG |
| R78K | F: AGTAGATTTCAAAGAACTTAATAAGAGAACTCAAGACTTCTG<br>R: AATTTTCTCCATTTAGTACTGTCTTTTTTCTTTATG |
| T290A | F: GAGGAACCAAAGCACTAGCAGAAGTAATACCACTAACAGAAG<br>R: TAGTGCTTTGGTTCCTCTAAGGAGTTTAC |
| T290D | F: GAGGAACCAAAGCACTAGATGAAGTAATACCACTAACAGAAG<br>R: TAGTGCTTTGGTTCCTCTAAGGAGTTTAC |
| T290S | F: GAGGAACCAAAGCACTAAGCGAAGTAATACCACTAACAGAAG<br>R: TAAGGAGTTTACATAATTGCCTTACTTTAATCCCTGGGTAAATC |

**Supplementary Table 2: Primers used in pRT6H-PROT site-directed mutagenesis.** The pRT6H-PROT-WT plasmid was used as the template to generate all single mutants. For double mutants, the specific template plasmid utilized is indicated in the table. Underlined nucleotides denote the introduced mutations

ApaI Forward: CATAGCCAAAAATTGCAGGGCC (forward primer for all fragment 1s)

SalI Reverse: GTAACGCCTATTCTGCTATGTCGA (reverse primer for all fragment 2s)

| Mutant | Primers (5' to 3') |
|---|---|
| W24A | Fragment 1 R: CTGTCAATGGCGCTTGTTTAACTTTTGGGCCATCC<br>Fragment 2 F: GTTAAACAAGCGCCATTGACAGAAGAAAAAATAAAAG |
| W24F | Fragment 1 R: CTGTCAATGGAAATTGTTTAACTTTTGGGCCATCC<br>Fragment 2 F: GTTAAACAATTTCCATTGACAGAAGAAAAAATAAAAG |
| F61A | Fragment 1 R: CTTTATGGCAGCTACTGGAGTATTGTATGGATTTTCAGGC<br>Fragment 2 F: CTCCAGTAGCTGCCATAAAGAAAAAAGACAGTACTAAATGG |
| F61L | Fragment 1 R: CTTTATGGCAAGTACTGGAGTATTGTATGGATTTTCAGGC<br>Fragment 2 F: CTCCAGTACTTGCCATAAAGAAAAAAGACAGTACTAAATGG |
| W24A/F61A<br>(from W24A) | Fragment 1 R: CTTTATGGCAGCTACTGGAGTATTGTATGGATTTTCAGGC<br>Fragment 2 F: CTCCAGTAGCTGCCATAAAGAAAAAAGACAGTACTAAATGG |
| W24A/F61L<br>(from W24A) | Fragment 1 R: CTTTATGGCAAGTACTGGAGTATTGTATGGATTTTCAGGC<br>Fragment 2 F: CTCCAGTACTTGCCATAAAGAAAAAAGACAGTACTAAATGG |
| W24F/F61A<br>(from W24F) | Fragment 1 R: CTTTATGGCAGCTACTGGAGTATTGTATGGATTTTCAGGC<br>Fragment 2 F: CTCCAGTAGCTGCCATAAAGAAAAAAGACAGTACTAAATGG |
| W24F/F61L<br>(from W24F) | Fragment 1 R: CTTTATGGCAAGTACTGGAGTATTGTATGGATTTTCAGGC<br>Fragment 2 F: CTCCAGTACTTGCCATAAAGAAAAAAGACAGTACTAAATGG |
| R78A | Fragment 1 R: GTTCTGCGAAATCTACTAATTTTCTCCATTTAGTACTGTC<br>Fragment 2 F: AGTAGATTTCGCAGAACTTAATAAGAGAACTCAAGATTTCTG |
| R78K | Fragment 1 R: GTTCTTTGAAATCTACTAATTTTCTCCATTTAGTACTGTC<br>Fragment 2 F: AGTAGATTTCAAAGAACTTAATAAGAGAACTCAAGATTTCTG |
| T290A | Fragment 1 R: CTACTTCTGCTAGTGCTTTGGTTCCCCTAAGAAG<br>Fragment 2 F: CCAAAGCACTAGCAGAAGTAGTACCACTAACAGAAGAAGC |
| T290D | Fragment 1 R: CTACTTCGTCTAGTGCTTTGGTTCCCCTAAGAAG<br>Fragment 2 F: CCAAAGCACTAGACGAAGTAGTACCACTAACAGAAGAAGC |
| T290S | Fragment 1 R: CTACTTCTGATAGTGCTTTGGTTCCCCTAAGAAG<br>Fragment 2 F: CCAAAGCACTATCAGAAGTAGTACCACTAACAGAAGAAGC |

**Supplementary Table 3: Primers used in pNL4-3ΔEnv-GFP mutagenesis.** The pNL4-3ΔEnv-GFP-WT plasmid was used as the template to generate all single mutants. For double mutants, the specific template plasmid utilized is indicated in the table. All mutagenesis reactions used a common forward primer containing an ApaI restriction site (for Fragment 1) and a common reverse primer containing a SalI restriction site (for Fragment 2). The mutation-specific internal primers for each fragment are shown in the table. Underlined nucleotides indicate sites of induced mutations.
