## Supplementary material for "Structural Basis of Polypurine Track Strand Displacement by HIV-1 Reverse Transcriptase": Descriptions for Supplementary Videos

**Description of Supplementary Videos for**

**Supplementary Video 1 | 3D variability analysis of the RT/T-P-$D_{PPT_{\underline{DNA}}}$/dATP complex.**

This video shows a series of cryo-EM maps (pink) derived from the RT/T-P-$D_{PPT_{\underline{DNA}}}$/dATP particle stack, capturing the flexibility of the downstream template-displacement duplex. A ribbon model of the consensus map containing 15 downstream nucleotides (blue) is fitted into a representative map in the series. **First close-up**: A zoomed-in view of the $RT_{p66}$ thumb subdomain and the displacement strand reveals that contact between T290 and the $PPT_{DNA}$ displacement strand occurs in only a subset of the maps, consistent with the 3D classification analysis. **Second and third close-ups**: A zoomed-in view of the $RT_{p66}$ fingers subdomain at the strand displacement (SD) site shows R78 contacting the template strand. Furthermore, F61 shifts to contact with the template strand, while W24 remains relatively fixed across all conformations. These observations collectively support the hypothesis that F61 and R78 coordinate template strand translocation, whereas W24 helps stabilize the displacement strand.

**Supplementary Video 2 | 3D variability analysis of the RT/T-P-$D_{PPT_{\underline{RNA}}}$/dATP complex.**

This video shows a series of cryo-EM maps (green) derived from the RT/T-P-$D_{PPT_{\underline{RNA}}}$/dATP particle stack, capturing the flexibility of the downstream template-displacement duplex. A ribbon model of the consensus map containing 17 downstream nucleotides (salmon pink) is fitted into a representative map in the series. **First close-up**: A zoomed-in view of the $RT_{p66}$ thumb subdomain and the displacement strand reveals that contact between T290 and the $PPT_{RNA}$ displacement strand occurs in only a subset of the maps, consistent with the 3D classification analysis. **Second and third close-ups**: A zoomed-in view of the $RT_{p66}$ fingers subdomain at the strand displacement (SD) site shows R78 contacting the template strand. Furthermore, F61 shifts to contact with the template strand, while W24 remains relatively fixed across all conformations. These observations collectively support the hypothesis that F61 and R78 coordinate template strand translocation, whereas W24 helps stabilize the displacement strand.
